# Promises and pitfalls of Topological Data Analysis for brain connectivity analysis

**DOI:** 10.1101/2021.02.10.430469

**Authors:** Luigi Caputi, Anna Pidnebesna, Jaroslav Hlinka

**Author notes:** Email addresses (Luigi Caputi), (Anna Pidnebesna), (Jaroslav Hlinka).

## Abstract

Developing sensitive and reliable methods to distinguish normal and abnormal brain states is a key neuroscientific challenge. Topological Data Analysis, despite its relative novelty, already generated many promising applications, including in neuroscience. We conjecture its prominent tool of persistent homology may benefit from going beyond analysing structural and functional connectivity to effective connectivity graphs capturing the direct causal interactions or information flows. Therefore, we assess the potential of persistent homology to directed brain network analysis by testing its discriminatory power in two distinctive examples of disease-related brain connectivity alterations: epilepsy and schizophrenia. We estimate connectivity from functional magnetic resonance imaging and electrophysiology data, employ Persistent Homology and quantify its ability to distinguish healthy from diseased brain states by applying a support vector machine to features quantifying persistent homology structure.

We show how this novel approach compares to classification using standard undirected approaches and original connectivity matrices. In the schizophrenia classification, topological data analysis generally performs close to random, while classifications from raw connectivity perform substantially better; potentially due to topographical, rather than topological, specificity of the differences. In the easier task of seizure discrimination from scalp electroencephalography data, classification based on persistent homology features generally reached comparable performance to using raw connectivity, albeit with typically smaller accuracies obtained for the directed (effective) connectivity compared to the undirected (functional) connectivity. Specific applications for topological data analysis may open when direct comparison of connectivity matrices is unsuitable - such as for intracranial electrophysiology with individual number and location of measurements. While standard homology performed overall better than directed homology, this could be due to notorious technical problems of accurate effective connectivity estimation.

## 1. Introduction

The general idea that mathematical approaches from geometry and algebraic topology can provide *powerful tools and methods for the understanding of complex data is fairly recent (see, e*.*g*., (Frosini, 1990; Edelsbrunner et al., 2002; Zomorodian and Carlsson, 2005; Carlsson, 2009)) and has lead to the birth and advancement of a completely new research branch, *Topological Data Analysis* (TDA). This is a fast growing and promising field at the intersection of Data Analysis, Algebraic Topology, Computational Geometry, Machine Learning and Statistics. Its main advantage is the possibility to analyse data by looking at its shape and underlying geometric structures arising from high-dimensional relations; this goal is achieved by providing a strong mathematical framework and manageable quantitative tools such as homology theory (Munkres, 1984; Hatcher, 2000). The structure of the data is then qualitatively and quantitatively assessed in the form of topological features (e.g. connected components, voids, tunnels or loops). This approach has been already widely applied and finds nowadays applications in many science domains dealing with various complex systems (Wasserman, 2018; Chazal and Michel, 2017).

The human brain and its dynamical activity represents one of the archetypal examples of complex systems. Indeed, due to both the intricate nature of the structure of connections of its constituent units, as well as the highly nonlinear and potentially chaotic dynamics within each of these units, it is rightly considered a suitable target example system for application of such complex system methodology. In particular, the study of the complex patterns of brain network topology has become a flourishing areas of research since the 2000’s (Bullmore and Sporns, 2009).

There are three main approaches to assess the interactions between brain regions: structural, functional, and effective connectivity (see Friston (1994, 2011); Park and Friston (2013)). Structural connectivity is the pattern of physical connections between the considered brain regions, while functional and effective are based on neuronal activity signals. Functional connectivity is defined as the stochastic dependence between remote neurophysiological events and provides thus a non-directional (symmetric) measure of brain interactions. On the contrary, the effective connectivity is defined as the direct influences that one region exerts over another. The most frequently used measure of functional connectivity is the (linear) Pearson correlation, which provides an accurate functional connectivity estimate for functional magnetic resonance imaging data under common processing settings (Hlinka et al., 2011), (Hartman et al., 2011). One of the applications in clinical neuroscience is the characterization of functional brain alterations in various diseases. The classification power of FC has been reported for schizophrenia (Lynall et al., 2010), multiple sclerosis (Zurita et al.), epilepsy (Liao et al., 2010), and many other brain diseases and states. On the other hand, effective connectivity, while in principle more informative with respect to the underlying dynamics since it specifies the direct influences among the brain regions, is more scarcely applied in practice, presumably since it is also much harder to estimate and the methods thus typically can handle only smaller networks (Smith, 2012). While there are many competing methods for estimation of effective connectivity offering different trade-off between advantages and drawbacks (Valdes-Sosa et al., 2011), there is a general agreement that if estimated well, they would provide valuable extra information on top of functional connectivity (Smith, 2012).

While a connectivity matrix characterizes only the strengths of all pair-wise interactions, more complex characterizations of the whole network as well as specific nodes thereof can be provided. This can be done by application of graph-theoretical tools to the graph constructed by using the connectivity matrix as an adjacency matrix (Telesford et al., 2011). However, the graph-theoretical analysis of brain connectivity suffers from multiple technical difficulties. One of them is that the classical approach requires applying a threshold to select which connections are relevant, i.e., strong enough to constitute an edge in the graph. There are multiple heuristics to deal with the threshold selection or even generalization of many graph metrics to weighted graphs, however no single optimal solution is available. Among other attempts Gallos et al. (2012), a very attractive way to side-step the problem and consider all possible thresholds at once, without losing any information, is thus offered by already established methods of TDA, such as Persistent Homology.

*Persistent Homology* (PH) is one of the main tools adopted in TDA and is a multi-scale adaptation of the classical homology theories studied in Algebraic Topology. It allows a computation of the *persistent* topological features of a space at all different resolutions (here: densities of the thresholded connectivity graph), while revealing the most essential ones. In the PH-approach to complex networks, instead of focusing on a single thresholded graph, one analyses a family of simplicial complexes indexed by a (filtration) parameter, by considering all the possible thresholds at once.

Based on a robust mathematical theory and being stable with respect to small noise perturbations (Cohen-Steiner et al., 2007), PH has provided a novel qualitative and quantitative tool to study complex data (in the form of either point clouds, time series or connectivity networks). Its use has already been reported in many different fields: in neuroscience and neurological disorders (Lee et al., 2011b), in endoscopy analysis (Dunaeva et al., 2016), angiography (Bendich et al., 2014), pulmonary diseases (Brodzki et al., 2018), finance (Gidea, 2017), fingerprint classification (Giansiracusa et al., 2019), image classification (Dey et al., 2017), to name few applications. The pipeline involving Persistent Homology in network analysis usually starts by considering (undirected, weighted) graphs as input. However, in many real world examples as in the study of effective connectivity, the input graphs are also equipped with a richer structure on the edges, namely the direction from the source node to the target. As directed graphs theoretically hold more information than their undirected counterparts (see e.g., Sec. 3.4), an extension of the PH pipeline to asymmetric networks might provide a more refined family of invariants. Some novel approaches have already been proposed in this direction by Masulli and Villa (2016); Reimann et al. (2017); Chowdhury and Mémoli (2018); Turner (2019); Aktas et al. (2019); Lütgehetmann et al. (2020), although not yet widely adopted; notably the early applications include the analysis of brain structural connectivity networks (Reimann et al., 2017).

The developments in the fields of brain connectivity analysis and TDA call for their combined utilization. Many attempts have been done in this direction, for reviews see (Solo et al., 2018; Phinyomark et al., 2017; Lee, 2019). In this work, with the aim of investigating the promise of applications of topological analysis to effective connectivity networks, we provide a pioneering example of such endeavour for the task of classifying altered brain states in several settings, while we assess the improvement provided by these methodologies in contrast to dealing with the connectivity directly, or applying the TDA approach to functional connectivity data only.

To this end, we construct functional and effective connectivity matrices (in short, FC and EC) derived from brain activity time series; these are obtained by two most common functional neuroimaging modalities – (intracranial) electroencephalography ((i)EEG) and functional magnetic resonance imaging (fMRI) – using two prominent examples of disconnectivity-related brain diseases: epilepsy and schizophrenia. For epilepsy, for a group of patients, we distinguish between ictal and pre-ictal connectivities. For schizophrenia, we classify patients from the matching neurotypical controls. Following previous approaches Lee et al. (2011b, 2012); Merelli et al. (2016); Stolz et al. (2018), we calculate the persistent homology of the (directed) filtrations associated to the brain functional (or effective) connectivity networks, and compute the associated PH-features (further referred to as *directed persistent homology*). In order to quantify their informative power, we apply a state-of-art classifier (a support vector machine, SVM), with the task to distinguish between healthy and diseased brain states. We use a range of most common quantitative PH features, namely *Persistence Landscapes* (PL) Bubenik and Dłotko (2017), *Persistence Images* (PI) Adams et al. (2017), *Persistent Entropy* (PE) Rucco et al. (2014) and *Carlsson Coordinates* (CC) Adcock et al. (2013). We compare the results with those obtained, under the same tasks, by a naive (SVM) classification applied to the FC and EC matrices directly.

The directed homology pipeline followed in this work was introduced in (Reimann et al., 2017) with applications in the study of the structural and functional connectivity in rat brain slices. This was also the first, and the only one known to the authors, application of this approach in neuroscience.

Note that while the application of directed persistent homology analysis to human brain effective connectivity is novel, it builds upon the tools and ideas developed and applied recently in the field. For example, the undirected variants of the TDA features have been shown to be promising in applications to Alzheimer (Kuang et al., 2019b; Pachauri et al., 2011), ASD (Wong et al., 2016), depression (Khalid et al., 2014), Parkinson (Garg et al., 2016), or in the study of childhood stress (Chung et al., 2014). The first work, applying PH-methods to brain networks constructed by using Pearson correlation is due to Lee (see (Lee et al., 2011b, 2012)) in the case of patients affected by ADHD. Among previous approaches in epilepsy, a persistent homology analysis has been already reported in (Merelli et al., 2016), where (weighted) PE is employed in order to detect the transition between the pre-ictal and ictal states, or in (Piangerelli et al., 2018; Wang et al., 2015), where a classification analysis in epilepsy, based on PLs and PE, has used different filtrations (sublevel-based or lower star filtrations). In (Choi et al., 2014), a study of the epileptic abnormality is given in rat models. Within schizophrenia applications, the only analysis applying methods from TDA is, to the authors knowledge, the work (Stolz et al., 2018). To provide more insight into the promises and pitfalls of the application of directed persistent homology analysis of brain (effective) connectivity, the current study goes beyond a single method application, and provides comparisons with the application of (standard) persistent homology of undirected networks, as well as with the application of machine learning directly to the input connectivity matrices. This, together with tackling three types of neuroimaging data across two brain different dysconnectivity conditions, namely through an easier classification task of ictal versus pre-ictal brain states, and a more compelling one, between schizophrenia patients and healthy controls, should provide a richer picture concerning the challenges of TDA application to brain connectivity data analysis.

Our key observation is that standard TDA applied to effective connectivity is in principle able to discriminate diseased from healthy brain states, comparably to (previously used combinations of) TDA applied to functional connectivity. However, in many practical situations such as we demonstrate in the results section for the application to schizophrenia classification, the use of raw functional connectivity features is advantageous. This may be due to topographical, rather than topological, specificity of the particular changes related to schizophrenia – these topographically (regionally/spatially) specific changes are naturally picked up by classifiers using the raw functional connectivity matrices; while the topological methods effectively ignore the spatial information, as they are invariant with respect to permutation of nodes – a fact that may be commonly overlooked when considering the applications. However, in yet another scenarios, the direct connectivity comparison is not straightforwardly applicable, such as for inter-subject analysis of intracranial electrophysiology recordings in epilepsy, not only due to potentially different spatial location of the brain alterations, but also due to the technical challenge of inter-individual differences in the number and location of the measurement sites.

With the idea that the topological pipeline may suffer from heterogeneous subject biases, a probable cause of lower general performances, we have also introduced an alternative methodological pipeline reducing the underlying heterogeneity. We observe that this variation of the pipeline shows a remarkable increment of the performances in the TDA-approach, when applied to the EEG dataset.

The paper is structured as follows. In Sec. 2 we recall the definition of directed Persistent Homology. In Sec. 3, we describe our pipeline and the common featurizations adopted in the classification tasks; we report our results in Sec. 4. In Sec. 5, we discuss our contributions. In the Appendix A, we review some mathematical concepts in more detail, including simplicial complexes, filtrations and (persistent) homology groups. Appendix B and Appendix C provide additional details concerning the data selection and processing. We conclude with some additional results (Appendix D) and a statistical comparison of PH versus DPH (Appendix F).

## 2. Persistent Homology of directed networks

In this section, following Reimann et al. (2017) and Lütgehetmann et al. (2020), we review the classical notions of simplicial complexes and persistent homology groups adapted to the *directed* framework, keeping always in mind the directed networks as the application example. The main objects in this context are directed graphs, and as in the context of undirected graphs, starting from a directed network we will construct suitable simplicial complexes, filtrations and persistent homology groups. We assume that the reader is familiar with these classical general concepts in the undirected case (reviewed for convenience in Appendix A); we refer to Zomorodian and Carlsson (2005) and to Otter et al. (2017) for an introduction of Persistent Homology with an eye towards applications.

A graph *G* = (*V, E*), where *V* denotes the set of its vertices and *E* the set of its edges, is a *directed graph* if every edge *e* = (*v*_0_, *v*_1_) is given by an *ordered* pair of *distinct* vertices *v*_0_ (the source of the edge) and *v*_1_ (the target). For a given pair of vertices *v*_0_ and *v*_1_, both edges (*v*_0_, *v*_1_) and (*v*_1_, *v*_0_) are allowed, but loops, i.e., edges of type (*v, v*), are not. A higher dimensional generalization of directed graphs is given by the notion of directed simplicial complexes (see Def. A.1 for the classical definition of simplicial complexes):

### Definition 2.1.

An *ordered simplicial complex* Σ = {*σ*_*α*_}_*α*_ on a vertex set *V* is a non-empty family of finite ordered subsets *σ*_*α*_ ⊆ *V* with the property that, if *σ* belongs to Σ then every ordered subset *τ* of *σ* (ordered with the natural order induced by *σ*) belongs to Σ.

In analogy with the notation used for simplicial complexes, the elements of an ordered simplicial complex of cardinality *d* + 1 are called *d*-*simplices*. We say that an ordered simplicial complex Σ has dimension *d* if all its simplices have cardinality at most *d* + 1. For example, every directed graph (*V, E*) is a 1-dimensional ordered simplicial complex on the set of vertices *V* and, if *V* is the set {0, …, *n*}, then the collection of all the ordered subsets of *V* is an *n*-dimensional ordered simplicial complex. When *n* = 1 and *V* = {0, 1}, the ordered simplicial complex given by the ordered subsets of *V* is nothing but the directed graph with vertices 0 and 1 and ordered edges (0, 1) and (1, 0).

In the context of undirected graphs, there are various ways to construct associated simplicial complexes; one of the most established ones is given by the clique complex, where one builds the simplicial complex associated to a graph *G* by using the combinatorial information of its cliques (see Ex. A.3). Following the analogy with this classical construction, from a given directed graph *G* we will construct an ordered simplicial complex by using the combinatorial information given by the ordered cliques of *G*, as we now explain. Recall that a *k*-clique in an unordered graph *G* is a complete subgraph on *k* vertices. Likewise, if *G* = (*V, E*) is a directed graph, then an *ordered k-clique* of *G* is a *k*-tuple (*v*_1_, …, *v*_*k*_) of vertices of *G* with the property that the pair (*v*_*i*_, *v*_*j*_) is an ordered edge of *G* for every *i < j*. We observe that in an ordered clique there is a source vertex, *v*_1_, having only out-edges, and a target vertex, *v*_*k*_, having only in-edges. For example, in the following graph on *V* = {0, 1, 2, 3}:

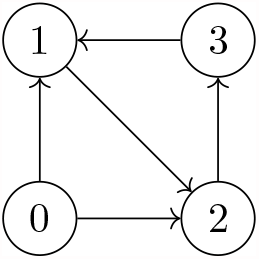

we have only one ordered 3-clique (in addition to the 1- and 2-cliques), *i*.*e*., the clique corresponding to the complete subgraph (0, 1, 2) on the vertices 0, 1 and 2. Observe that the subgraph on the vertices 1, 2 and 3, although being a 3-clique for the underlying undirected graph, is not an *ordered* 3-clique as there is no common source or target (it is, in fact, a cycle). For a given directed graph, we can construct an ordered simplicial complex by considering the collection of all the directed cliques (see the analogous construction of Ex. A.3):

### Definition 2.2.

The *directed flag complex* dFl(*G*) associated to a directed graph *G* is the ordered simplicial complex whose *k*-simplices are all the ordered (*k* + 1)-cliques of *G*.

Observe that the vertices and edges of a directed graph *G* are exactly the 1- and 2-cliques of *G*, and that every directed graph is contained in its associated directed flag complex. For example, if *G* is the directed graph on *V* = {0, 1, 2}, with edges (0, 1), (1, 2) and (0, 2), then there is only one directed 3-clique, (0, 1, 2), and the associated directed flag complex dFl(*G*) consists of the same vertices and edges of *G*, together with the 2-simplex (0, 1, 2) corresponding to the unique directed clique:

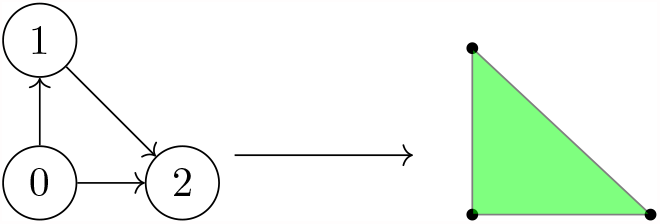

In order to study persistent homology groups, once we have a suitable notion of simplicial complexes, we need filtrations of simplicial complexes. A filtration of ordered simplicial complexes consists of a nested sequence of increasing ordered simplicial complexes (see also Def. A.4). Various filtrations are generally possible. In Ex. A.5 we review the clique filtration and in Ex. A.6 the Rips filtration, an instance of the clique filtration (see Remark A.7) for undirected graphs. In the context of directed graphs, we construct filtrations using the directed flag complexes, as the natural extension of the clique filtration to the directed context. To be more precise, if Ø ⊆*G*_0_ ⊆ … ⊆ *G*_*n*_ = *G* is a family of directed subgraphs of a directed graph *G*, then the sequence

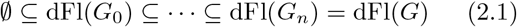

of directed flag complexes dFl(*G*_*i*_) is a filtration of dFl(*G*).

In our applications, we first consider a *weighted* directed graph *G*, i.e., a directed graph for which every edge is given a numerical weight. For each weight *w*_*i*_ we define the directed graph *G*_*i*_ as the (directed) subgraph of *G* with same vertices of *G* and only those edges corresponding to the weights *w* of *G* with *w* ≤ *w*_*i*_ (note that at this level the weights correspond already to a distance, rather than similarity matrix, see also Ex. A.5). The weights *w*_*i*_ of *G*, sorted in the increasing order, induce a sequence of directed subgraphs of *G*, hence a filtration of directed flag complexes dFl(*G*_*i*_). Generally speaking, in order to extend it to a filtration of directed flag complexes one has to choose a weighting function also on the directed simplices (hence on the ordered cliques) extending the weights given on the edges. We use the intuitive maximum weighting function (the default in the software *Flagser* Lütgehetmann et al. (2020)), associating to an ordered simplex the maximal weight across all its edges. See Lütgehetmann et al. (2020) or the documentation at https://github.com/luetge/flagser for details on other possible weighting functions.

Once a filtration of (ordered) simplicial complexes is given, one can apply the classical homology methods for obtaining Persistent Homology groups, barcodes and Persistence Diagrams (see App. A.2 for a review). We will briefly refer to the Persistent Homology of the filtration of directed flag complexes associated to a weighted directed graph as its *directed Persistent Homology* (shortly denoted by DPH).

## 3. Methods

In the analysis of brain connectivity networks, we use Persistent Homology features as input for classification by an SVM classifier. The steps of the pipeline include (see also Figure 1):

**Figure 1:**
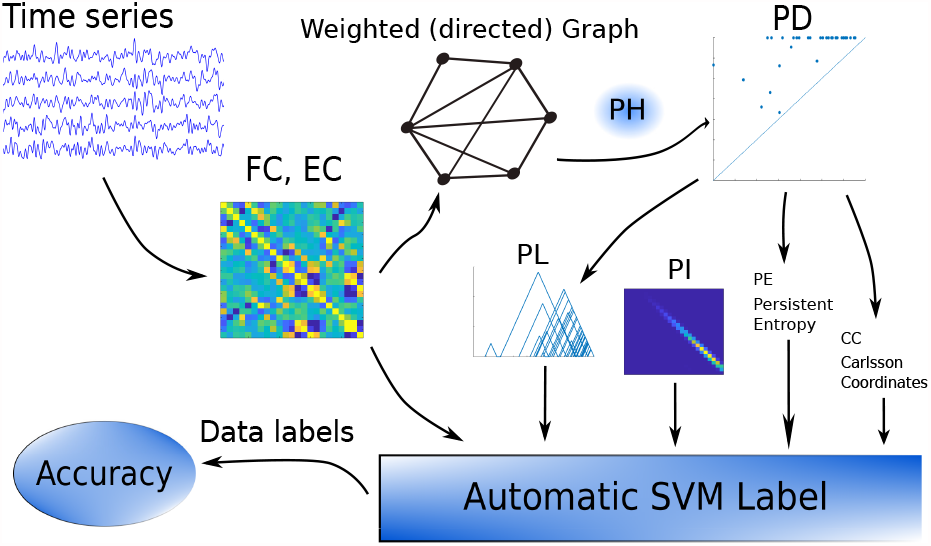
Pipeline of the main analysis. From the time series, we estimate the connectivity matrix, based on which build a weighted graph. After estimating Persistent Homology (PH) of the graph, we get Persistent Diagram (PD), that is represented by different features (Persistent Landscape PL, Persistent Image PI, Persistent Entropy PE, Carlsson Coordinates CC) for classification. Finally, classification based on PH features is compared with classification using matrices itself.

### Data

we use scalp and intracranial EEG (of epileptic patients) and fMRI (of schizophrenia patients and healthy controls) data in the form of multi-variate time series;

### Graphs

for each dataset we construct (functional and effective) connectivity graphs by calculating Pearson correlation and Granger Causality respectively; see Sec. 3.2.

### Topology

from the graphs we construct appropriate filtrations of simplicial complexes (see Sec. A.1 and Sec. 2). In particular, we have constructed the Vietoris-Rips filtration (see Ex. A.6) associated to the FC graph and the filtration of directed flag complexes associated to the directed EC graph (see Eq. (2.1)).

### Homology

we compute the Persistent Homology of the given filtrations (PH for undirected and DPH for directed graph), getting their persistence Betti numbers (corresponding to the number of connected components for dimension 0 and to the number of loops for dimension 1, the two most common settings in the literature); see also App. A.2.

### Machine Learning

from persistence Betti numbers, we get the features (vectors of real numbers) and apply a suitable classifier (SVM) to both the FC or EC graphs, and to the features (in particular, the Persistence Landscapes, Persistent Images, Carlsson Coordinates and Persistent Entropy (Sec. 3.5).

As a variation of this pipeline, we add an additional cleaning step in the case of EEG and iEEG datasets as described in Sec. 3.6.

#### 3.1. Dataset

We use two prominent types of neuroimaging data in combination with two examples of pathological brain dynamics:

##### 3.1.1. fMRI data (Schizophrenia)

We analyse the dataset that consists of fMRI recordings of 90 patients with a schizophrenia diagnosis and 90 healthy controls. Functional MRI data were collected with a 3T MR scanner (Siemens; Magnetom Trio) at Institute of Clinical and Experimental Medicine in Prague. Participants were informed about the experimental procedures and provided written informed consent. The study design was approved by the local Ethics Committee of the Institute of Clinical and Experimental Medicine and the Psychiatric Center Prague. *T* 2^∗^-weighted images with BOLD contrast were acquired with voxel size 3×3×3mm^3^ and TR/TE parameters of 2000/30ms. As anatomical reference the T1-weighted high-resolution image was used (TR/TE/TI = 2300/4.6/900 ms, voxel size = 1×1×1mm^3^). Initial data preprocessing was done using FSL routines (FMRIB Software Library v5.0, Analysis Group, FMRIB, Oxford, UK) and CONN toolbox (McGovern Institute for Brain Research, MIT, USA). Functional realignment and unwarping, slice-timing correction, structural segmentation into white matter and cerebrospinal fluid and structural normalization to the MNI space were done using CONN’s default preprocessing pipeline (defaultMNI), as well as functional normalization to the MNI space, outlier detection, and smoothing with 8mm kernel size. The estimated amount of motion in the subjects was generally satisfactory, with the average framewise displacement between subsequent volumes being 0.13 ± 0.08mm (mean ± std across subjects). Six head-motion parameters with their first order temporal derivatives and five principal components of white-matter and cerebrospinal fluid time-series were regressed out to suppress measurement artifacts. Furthermore, to remove possible signal drift, time-series were linearly detrended and filtered by band-pass filter [0.009-0.08Hz]. See Kopal et al. (2020) and Oliver et al. (2019) for detailed prepocessing description. To extract the time series for further analysis, the brain’s spatial domain was divided into 90 non-overlapping regions of interest (ROIs) according to the AAL atlas; from each ROI we extract one BOLD time series by averaging the time series of all voxels in the ROI.

##### 3.1.2. EEG data (Epilepsy)

We analyse both scalp and intracranial electroencephalography EEG data:

### Scalp EEG data

We use EEG recordings from pediatric subjects with intractable seizures, collected at the Children’s Hospital Boston. The data are available at https://physionet.org/content/chbmit/1.0.0/, from the publicly open dataset *Physionet*. We analyse EEGs belonging to 18 subjects, with 23 scalp electrodes. In particular, we included patients with typical epileptic event of length between 20 and 120 seconds. The set of selected patients is provided in Appendix B. After isolating the epileptic event according to labels in the patient’s summary file, we also consider non-epileptic records, chosen randomly from the database (we extract segments of the same length as the seizure segments). In total, we use 102 ictal and 102 interictal data segments, the same amount of ictal and preictal parts for every particular subject. Sampling frequency is 256Hz, no special preprocessing is applied except for a band-pass filter [1-70Hz], notch filter at 50Hz, and removal of global signal for every data segment separately.

Several seizure recordings are available for each subject. Therefore, to avoid dependence between samples and any potential biases due to different numbers of seizure recordings available, for each subject we average all the connectivity matrices for ictal (interictal) periods. Thus, the dataset for the analysis consists of two connectivity matrices per subject (one ictal and one inter-ictal). For the additional analysis using a modified pipeline (Section 3.6) we used a slightly modified data selection, which is described in Appendix B.

### Intracranial EEG data

We use the anonymized dataset of 16 patients of the epilepsy surgery program of the Inselspital Bern, freely available at http://ieeg-swez.ethz.ch/, with no further preprocessing implemented. The intracranial electroencephalography (iEEG) recordings belong to 16 patients with pharmacoresistant epilepsies collected at the Sleep-Wake-Epilepsy-Center (SWEC) of the University Department of Neurology at the *Inselspital Bern* and the *Integrated Systems Laboratory* of the ETH Zurich. Each recording consists of 3 minutes of preictal segments, the ictal segment (ranging from 10 s to 1002 s), and 3 minutes of postictal time. In this work we only consider the recordings corresponding to the first 30 seconds of the first seizure as the ‘ictal’ data, and the 30 seconds segment starting 1 min before the first seizure start as our ‘preictal’ data.

For every patient we use the first minute of interictal data for estimation of baseline connectivity, which is later subtracted from both ictal and preictal connectivities.

#### 3.2. Connectivity Networks

In order to estimate the brain networks we use both a symmetric (Functional Connectivity, Sec. 3.2.1) and an asymmetric approach (Effective Connectivity, Sec. 3.2.2):

##### 3.2.1. Functional Connectivity

To obtain the functional connectivity network, we cross-correlate the time-series described in Sec. 3.1.2 and Sec. 3.1.1. Let *N* be the number of brain regions in the AAL atlas or number of electrodes in scalp EEG, then the *N* × *N* matrix *A* = (*A*_*i,j*_) is defined using Pearson correlation between the time series corresponding to nodes *i* and *j*. In line with previous literature (Lee et al., 2011b, Section 3), (Merelli et al., 2016, Section 3), (Stolz et al., 2018, Section 2.2), all entries of *A* that are not significant at *α* = 0.05 are set to 0, along with all negative values of functional connectivity. Below, we refer to this method as *masked Functional Connectivity (masked FC)*.

##### 3.2.2. Effective connectivity

The effective connectivity is quantified by the pairwise Granger causality (Barnett and Seth, 2013); furthermore, edges corresponding to links that do not show statistically significant causal effect (by means of the FACDA algorithm (Kořenek and Hlinka, 2020)) have their weight set to zero. In the following we refer to this method as *masked Effective Connectivity*, or more concisely, as *masked EC*. We use pairwise Granger causality instead of multivariate Granger Causality (which controls for all the other variables) to reach more robust estimates. Indeed, in the multivariate approach a greater number of parameters has to be estimated, potentially leading to less accurate estimates (Hlinka et al., 2013).

#### 3.3. Persistent Homology computation

We analyse persistent homology groups of both the directed and undirected networks. We refer to Appendix A for a review of PH. When not otherwise specified, the persistent homology computations shown in the article refer to calculations implemented in dimension 1, that is detection 1-dimensional holes; we refer to Appendix D for the corresponding results using the dimension 0 features, corresponding to the detection of connected components); homology groups are computed over the commonly used base field 𝔽_2_ (see Def. A.8).

##### 3.3.1. PH of undirected networks

The undirected network constructed in Sec. 3.2 is a weighted graph. As explained in Ex. A.5, by sorting the weights in an increasing or decreasing order, we get a filtration of simplicial complexes, the WRCF. Equivalently, one can construct the associated Vietoris-Rips filtrations (see Ex. A.6). As the implementation of the WRCF can be computationally demanding, in this work we follow the second approach: we construct the quasi-metric space (*X, d*) associated to the matrix *A* (the matrix *A* being constructed as described in the previous subsection) with underlying space *X* given by the vertices of the weighted graph and quasi-distance *d*(*i, j*) := 1 − *A*(*i, j*). We then construct the Vietoris-Rips filtration associated to this metric space and compute the associated PH and persistence diagrams. The computation of the associated Persistent Homology has then been implemented with the python library *Ripser* (Tralie et al., 2018).

##### 3.3.2. PH of directed networks

The directed network presented in Sec. 3.2.2 provides a weighted directed graph, hence a directed flag complex as explained in Sec. 2. For consistency with the undirected case, we first rescale the matrix *A* via a linear map *A* ⟼ *Ã* to span the interval [0, 1] and then apply the transformation *x* ⟼ 1 − *x* to use the matrix 1 − *Ã*, getting a positive matrix where the lowest values represent the strongest connections. We consider the sequence of directed graphs sorted with the increasing order of the weights (of 1 − *Ã*). We extend the weights to the simplices of the induced flag complexes by giving to a simplex *σ* the maximum value across the weights of its edges. We then compute the associated persistent homology groups.

#### 3.4. Why DPH? A motivational example with VAR processes

In this subsection we elucidate why an extension of the persistent homology techniques to direct networks is theoretically more fruitful than the classical PH approach on undirected networks. We assume that the reader is familiar with the concept of VAR(*p*) (vector autoregressive) processes (see Lutkepohl (2007) for a general overview).

Let *X* be a VAR(1) process of the form

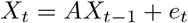

where *X*_*t*_ is an *n*-dimensional random vector, *A* is a *n* × *n* matrix of (real) coefficients and *e*_*t*_ is a white noise with 0 mean and covariance Σ; we say that the process *X* is *generated* by the matrix *A*.

Without loss of generality, we assume that *X* is *stable* (meaning that the matrix *A* has eigenvalues smaller in module than 1), has 0 mean and the variance Σ is the identity matrix Σ = 1. Then, the matrix sequence {*A*^*i*^} _*i*∈ℕ_ is absolutely summable and the covariance matrix of *X* (that is a stable process, hence stationary) can be written as

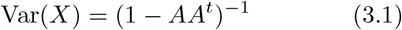

where *A*^*t*^ denotes the transpose of the matrix *A*.

The correlation matrix of the process *X* depends only on its covariance matrix Var(*X*) = Cov(*X, X*), whereas the associated effective connectivity depends upon the generating matrix *A* (for simplicity, we use the matrix A directly here as the EC).

In order to document the extra richness of the directed version of PH, we consider two different VAR(1) processes *X* and *Y* generated by two matrices *A* and *B* with different topological structure but the same covariance matrix Cov(X, X) = Cov(Y, Y). These two processes share the same correlation matrix (used in the definition of FC), hence the associated (standard/undirected) PH will not distinguish them. However, as the associated EC matrices differ, the *directed* PH might be able to distinguish the aforementioned processes. We now document this using a simple low dimensional example.

The problem reduces to finding topologically different matrices *A, B*, such that *AA*^*t*^ = *BB*^*t*^ (this can be achieved by choosing an suitable orthogonal matrix *Q*, and by setting *B* := *AQ*. As a specific example, we set:

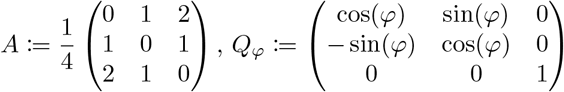

Let *φ* = *π/*3 and *B* = *AQ*_*φ*_. The eigenvalues of *A* and *B* are smaller in module than 1 and the generated VAR(1) processes are stable, moreover, clearly *AA*^*t*^ = *BB*^*t*^, hence, they have the same covariance matrix by Eq. (3.1), and as a consequence, the correlation matrices of *X* and *Y* have same persistent homology groups (see Fig. 2). On the other hand, when applied to the generating matrices *A* and *B*, DPH is able to distinguish the two processes (see Fig. 3), proving the advantage of DPH in distinguishing a larger class of processes.

**Figure 2:**
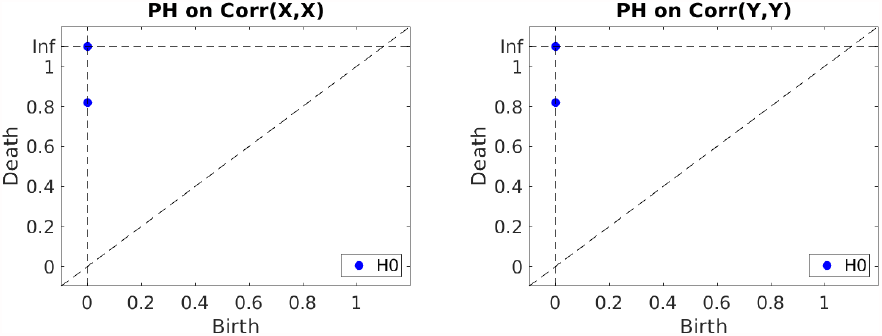
Left: PH of correlation matrix *Corr*(*X, X*). Right: PH of correlation matrix *Corr*(*Y, Y*).

**Figure 3:**
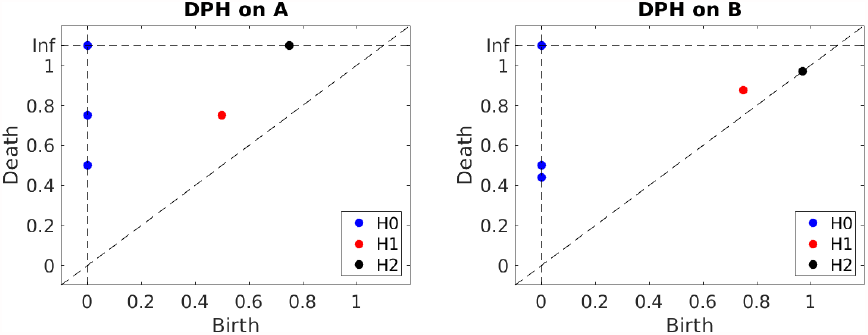
Left: DPH of the matrix *A*. Right: DPH of the matrix *B*.

We point out here that we cannot apply PH directly on *B*, as *B* is an asymmetric matrix. However, PH could indeed be in principle applied to the matrix *A* that is symmetric; however, it provides a different homology structure than the application of DPH. In particular, the persistent homology groups associated to *A* are concentrated in dimension 0, whereas DPH(A) has non-trivial homology classes in dimension 1 and 2 as well as in dimension 0 (see Fig. 4).

**Figure 4:**
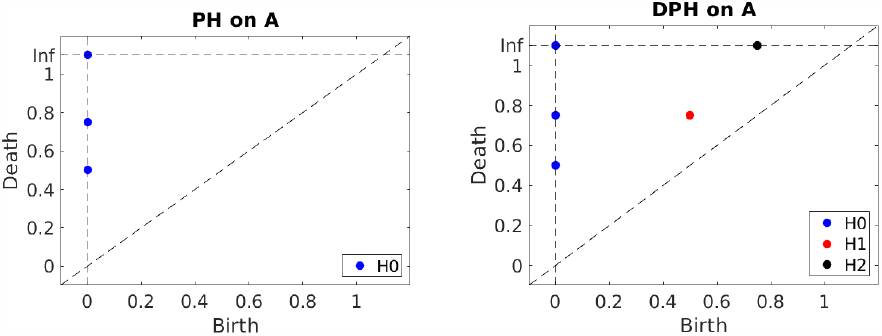
Left: PH of the matrix *A*. Right: DPH of the matrix *A*.

#### 3.5. Featurizations of Persistent Homology

Persistent homology groups can be visualized by using Persistence Diagrams as birth-death planar representations (Sec. A.2). In order to apply a SVM classifier, we derive four most commonly used vectorizations, or featurizations, from the persistence diagrams: Persistence Landscapes, Persistence Images, Carlsson Coordinates and Persistent Entropy. In this way we get input real-valued vectors amenable to the classifier. We proceed by illustrating the definitions of these topological features.

##### 3.5.1. Persistence Landscapes (PL)

*Persistence landscapes* have been introduced in order to perform PH-based statistical analysis by Bubenik and Dłotko (2017). For a birth-death interval (*b, d*), where *d* is a finite value, the function *f*_(*b,d*)_: ℝ→ ℝ *t* ↦ max(min(*t* − *b, d* − *t*), 0) represents a triangle-shaped function over (*b, d*). Let {(*b*_*i*_, *d*_*i*_)}_*i*∈*I*_ be a (finite) collection of birth-death intervals. Let *k*-max({*a*_*i*_}_*i*_) denote the *k*-th maximum value of a finite family of real numbers {*a*_*i*_}_*i*_.

###### Definition 3.1.

The *persistence landscape* of the set of birth-death coordinates {(*b*_*i*_, *d*_*i*_)}_*i*∈*I*_ is the family of functions *λ* = (*λ*_*k*_: ℝ → ℝ)_*k*∈ ℕ_, where 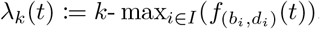.

Observe that the functions *λ*_*k*_(*t*) are piece-wise linear and that they are identically 0 when *k* is bigger than the cardinality |*I*| of the birth-death pairs. See Fig. 5 for an example of persistence landscape associated to a persistence diagram.

**Figure 5:**
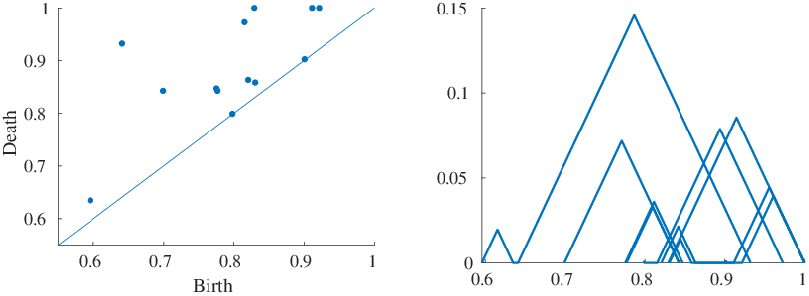
Left: persistence diagram (sample EEG data). Right: graphical representation of the related persistence landscape.

We follow the same method as Yesilli et al. (2019) to perform ML analysis on the set of persistence landscapes. Let 𝒟 = (*D*_*p*_)_*p*∈*P*_ be a set of persistence diagrams (*e.g*, each diagram *D*_*p*_ is the PD associated to a single brain network, at a fixed dimension). For every such persistence diagram *D*_*p*_, let 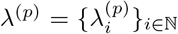be the associated PL. Let *K* ⊆ ℕ be a fixed subset of indices of ℕ; for every *k* ∈ *K* consider the collection of functions 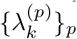 varying with the index *p* (and fixed *k*).

By definition, for every *k* and *p*, the map 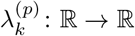 is a piecewise linear function. Then, it has a set of critical points 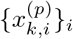, i.e., those points at which the function 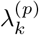 changes its direction. Let *z*^(*k*)^ be the vector of ℝ^*n*^ whose entries consist of all the critical points 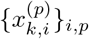 associated to the functions 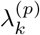, at *k* fixed; the entries of *z*^(*k*)^ are sorted in increasing order, without repetitions.

For every chosen *k* ∈ *K*, we have constructed a vector *z*^(*k*)^ depending only on the critical points of the *k*-th functions of the PLs. The feature vector, input of the classifier, associated to the diagram *D*_*p*_ in degree *k*, is then the evaluation vector 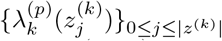. See the work of Chazal and Michel (2017) for further details. In our work, we choose the set *K* ⊆ *ℕ* to be *K* = {1} to capture the landscape containing information about the most persistent holes; hence, the analysis reported in Sec. 4 uses the first landscapes 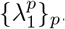.

##### 3.5.2. Persistence Images (PI)

An efficient and stable vector representation of persistent homology is given by the so called *Persistence Images* introduced by Adams et al. (2017). We start with a persistence diagram *D* = {(*b*_*i*_, *d*_*i*_)}_*i*∈*I*_, a subset of ℝ^2^ as depicted in the example Fig. A.8 and apply the transformation *T*: ℝ^2^ → ℝ^2^ sending a birth-death pair (*b*_*i*_, *d*_*i*_) to the birth-persistence pair (*b*_*i*_, *p*_*i*_) := (*b*_*i*_, *d*_*i*_ −*b*_*i*_). Let *D*_*k*_: ℝ^2^ → ℝ be the normalized gaussian centered at (*b*_*k*_, *p*_*k*_) and with standard deviation *σ*, i.e.,

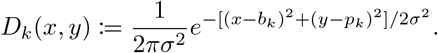

As set by default in line with Adams et al. (2017), we choose a specific non-negative weighting function *W*: ℝ^2^ → ℝthat is zero along the horizontal axis, continuous, and piece-wise differentiable, in particular a function depending only on the second coordinate: if *p* is the maximal possible persistence, *W* (*x, y*) is set to 0 if *y* ≤0, 1 if *y* ≥ *p* and *y/p* between 0 and 1.

We associate to the persistent diagram *D* a function *ρ*: ℝ^2^ → ℝdefined as weighted combination of gaussians:

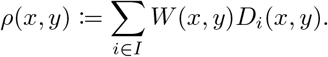

Let *b*_max_ and *p*_max_ be the maximum birth and maximum persistence across the births and persistences appearing in a diagram, i.e., *b*_max_ := max_*i*_(*b*_*i*_) and *p*_max_ := max_*i*_(*d*_*i*_ − *b*_*i*_). After choosing a grid of the domain [0, *b*_max_] × [0, *p*_max_], we compute the integral *∫ ρ*(*x, y*)*dxdy* at each pixel-box of the grid-domain. This gives a discretization of the surface image of *ρ* (ℝ^2^), hence a matrix of values which is our persistence image. See Fig. 6 for an example of the output.

**Figure 6:**
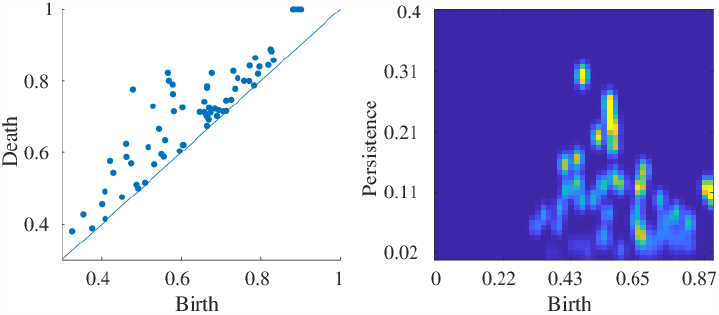
Left: Persistence diagram of example subject. Right: the associated PI, with domain grid [0, *b*_max_] *×* [0, *p*_max_].

For a family of persistence diagrams, one can define a *mean PI* of the persistence images associated to the PDs in the family. As each single PI is a matrix of real values, the mean PI of the family is defined as the matrix given by the element-wise average of the matrices. However, this brings a technical issue, as we describe:

###### Remark 3.2.

To construct the PIs, we have used the MATLAB code that accompanies the paper by Adams et al. (2017). Given a Persistence Diagram, one has to set parameters such as the resolution, the probability density and the weighting function. However, as already observed in (Stolz et al., 2018, Section 3.3), in order to compute the mean PI for a family of persistence diagrams, there is an additional pair of values to be chosen: the pair of maximum birth and maximum persistence. We set these values to the maximum across all subjects to achieve unbiased classification results. Note that an alternative group-specific normalization was reported to provide better classification results (Stolz et al., 2018), but in fact provides artificially inflated accuracy due to introducing group-specific bias; a topic that would be covered in more detail in a separate methodological report.

##### 3.5.3. Carlsson coordinates (CC)

Another useful featurization method is given by the so-called *Carlsson Coordinates* (Adcock et al., 2013, Sec. 4.1.2 & 4.2.2). The method is based on an approach from algebraic geometry. For every persistence diagram D = {(*b*_*i*_, *d*_*i*_)} _*i*∈*I*_ in the form of birth-death coordinates, we construct five vector features as follows. Let *p*_*i*_ = *d*_*i*_ − *b*_*i*_ be the life associated to (*b*_*i*_, *d*_*i*_) and let *p*_*M*_ be the maximal death time *d*_*M*_ := max_*i*_(*d*_*i*_). Following Adcock et al. (2013), we consider the following coordinates:

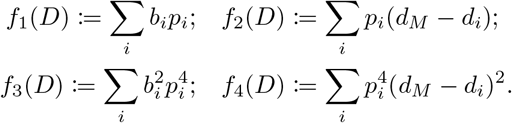

Morover, following (Yesilli et al., 2019, Sec. 4.4), we add a fifth coordinate *f*_5_(*D*) := max_*i*_(*p*_*i*_) taking into account the maximal reached persistence. As suggested in (Adcock et al., 2013, Sec. 4.2.2), we normalize the coordinates *f*_1_, …, *f*_4_ by a scale factor 1*/N*, where *N* is the number of the birth-death pairs in the persistence diagram.

##### 3.5.4. Persistent Entropy (PE)

Our last feature is *Persistent Entropy*, a topological adaptation of Shannon entropy in the context of persistent homology (Rucco et al., 2014), i.e., a number that measures the information encoded in the persistent homology groups (at a given dimension).

Let ℱ be a filtration of a simplicial complex Σ (see Def. A.4) and *D* = {(*b*_*i*_, *d*_*i*_)} _*i*∈*I*_ the set of birth-death coordinates associated to PH in dimension *k*. For every *i* ∈ *I* let *p*_*i*_ := *d*_*i*_ − *b*_*i*_ be the *i*-th persistence.

###### Definition 3.3.

The *persistent entropy E*(*B*) associated to *D* is the number *E*(*D*) := — ∑_*i*∈*I*_ *s*_*i*_ log(*s*_*i*_) where *s*_*i*_ = *p*_*i*_*/S*_*D*_ and *S*_*D*_ =∑ _*i*∈*I*_ *p*_*i*_.

#### 3.6. Controlling for inter-individual variability in electrode placement

In real-world scenarios, a substantial variability in the estimated functional or effective connectivity networks between subjects can be due to confounding purely anatomical differences, or spatial mis-alignment of the observed locations. In an experimental situation with repeated measurements from the same subject, the design may potentially allow suppressing this inter-individual variability by data preprocessing. In particular in iEEG data, a problem of data inconsistency across subjects arises, as every patient has his/her own unique position of electrodes (the implantation) that depends on the particular clinical requirement. Such implantation defines the basic structure of the connections between the recorded brain regions and, as a consequence, iEEG connectivity matrices can not even be directly compared between subjects because of the different data structure and the different sizes.

While this problem is much alleviated in scalp EEG with standardized electrode number and placement, inter-individual differences in brain and head anatomy may still cause some inter-individual variability in EEG connectivity that may obscure intra-individual variability reflecting the brain state dynamics of interest.

Working with the basic topological features rather than with the matrix directly should help to sidestep this problem. However this may not perfectly resolve the issue, as the geometry of the patient’s data may still be systematically biased by the individual structure of electrode implantation and anatomy. Therefore, before performing the classification task, we add two steps of data cleaning to the general pipeline.

The first step - used to decrease the influence of implantation on the topology - is directly applied to the connectivity matrices computed by the procedure described in Sec. 3.2. The EEG and iEEG data contain epileptic (ictal) and non-epileptic (preictal and interictal) segments as described in Sec. 3.1.2. In both cases, apart from the ictal and immediately preictal segments, we estimate the connectivity within the clearly interictal segments, which in the following we refer to as the *baseline connectivity*. This baseline connectivity is subtracted from both the preictal and ictal connectivities to provide correction for the confounding inter-individual variability.

The second step in the data cleaning aimed to further control for inter-subject variability is performed after the feature computation and before applying the classification tasks. For each topological feature and patient, we consider the vector consisting of all the realizations of the topological feature across conditions (i.e. all the classified segments for that subject). We compute the mean and remove it from the whole vector of feature values: this gives a final vector with null mean for every patient and feature that has inter-individual biases accounted for, and can be used in the classification tasks.

#### 3.7. Classification

To perform the classification (based on a set of features for each subject), we apply a linear support vector machine (SVM) that constructs a hyperplane in high-dimensional space on a training set of points. This hyperplane is then used for the classification of testing points.

We train a SVM classifier based on the topological features of Sec. 3.5. We have chosen the SVM as it is the state-of-art method when deep learning can not be applied due to insufficient sample size. Indeed, other models and classifiers could have been used, for example, a random forest approach instead of the SVM. A comparison of these methods employed in classification tasks on persistent homology of FC (albeit in autism rather than schizophrenia) has been recently provided by Rathore et al. (2019), where the methods have shown to achieve similar performances.

Evaluation of the classifier accuracy is made with the leave-two-out procedure on paired samples. In the following, when dealing with the epileptic datasets, by a “data point” we mean (the functional/effective) connectivity matrix characterizing particular brain activity (ictal or inter-ictal); when working with the schizophrenia dataset, “data points” are the single subject connectivities. For each dataset, we train the classifier on the set consisting of all the data points excluding both the tested sample and its matching counterpart. A similar approach has been used, for instance, in the studies by Castro et al. (2011), Mikolas et al. (2018) and Rubin-Falcone et al. (2018). To be more precise, for the epileptic datasets, for every tested connectivity, we exclude its match from the opposite type of activity (ictal/inter-ictal). For the schizophrenia data, for every tested subject, one subject from the opposite group was excluded, in order to keep groups of the same size. For all datasets, presented in the main text, such across-participants leave-two-out procedure was performed. In the supplementary, we also present analysis of EEG data, where classification was performed within-subject in selected subjects with high number of measured seizures, here always a matched pair of ictal and inter-ictal segment from the same recording is removed.

#### 3.8. Software

In the construction of effective connectivity networks we have used the software companion of Barnett and Seth (2013) for the computation of Granger Causality and of Kořenek and Hlinka (2020) for the construction of the mask (see Sec. 3.2.2). For the calculations of persistent homology groups of undirected networks we have used the python library *Ripser* Tralie et al. (2018). *For the computation of persistent homology of the directed graphs, we have used the library Flagser* by Lütgehetmann et al. (2020) (via the python API reference *pyflagser* of the Giotto-tda Tauzin et al. (2020), version v 0.2.0.), with the default settings. For computing the PIs, we have used the software that accompanies the paper of Adams et al. (2017); we set the variance of the Gaussians to 0.0001 and keep all the other parameters as by default. The pair maximum birth/maximum persistence is computed across all the instances (see Remark 3.2). For the calculation of PLs, we use our own MATLAB implementation of (Bubenik and Dłotko, 2017, Algorithm 1). For the implementation of support vector machine classifier, we use the MATLAB function *fitcsvm*. When otherwise specified, we use the default settings of the declared functions.

## 4. Results

The aim of the paper is to assess the applicability of (directed) persistent homology methods to brain functional/effective connectivity analysis. We have focused on two prominent neurological applications: epileptic seizure detection (using EEG data, both scalp and intracranial) and classification of schizophrenia/healthy brain states (using fMRI data). We follow the pipeline described in Sec. 3: for each subject, we first calculate the functional connectivity graph (Sec. 3.2.1) and the effective connectivity directed graph (Sec. 3.2.2), hence the associated Vietoris-Rips filtration (Ex. A.6) and the filtration of directed Flag complex (Def. 2.2), respectively; we then compute the persistent homology groups of the obtained filtrations as described in Sec. 3.3. We encode the PH information in the form of Persistence Diagrams, that we use to compute the associated topological features PL, PI, CC and PE. Here we potentially substract individual feature mean to control for electrode placement bias as described in Sec. 3.6 (this is not applicable to the fMRI data where the classification is only between subjects). We finally apply the SVM classifier (see Sec. 3.7). We compare the obtained classification results with those achieved by using the same SVM classification method applied to Functional Connectivity (FC) (in the undirected case) and to (masked) Effective Connectivity (EC) (in the directed approach). The results are shown in the subsections below. Note that for the results obtained on the iEEG data, a direct comparison with the naive FC/EC approach is not possible, because of different size and position of patient’s implantations.

We proceed by illustrating our results in the group analysis of the fMRI (schizophrenia) data first, of the group analysis of scalp EEG (epilepsy) data afterwards (following the classical pipeline first, and the alternative pipeline later), and we conclude with the analysis of the iEEG (epilepsy) dataset. The persistent homology computations shown in the following subsections refer to calculations in dimension 1 and we refer to Appendix D for the corresponding results in dimension 0.

### 4.1. First dataset: fMRI - Schizophrenia

In this first analysis, we present the results of a classification using the fMRI time series of 90 healthy subjects and of 90 schizophrenia patients (see Sec. 3.1.1). We report the results in Table 1.

**Table 1:** Schizophrenia: group analysis, 90 patients plus 90 healthy controls. SVM classification results using FC, EC and the topological features PL, PI, CC, PE in dimension 1 with undirected approach (PH) or directed approach (DPH). *Acc* stands for *Accuracy, Sns* for *Sensitivity* and *Spc* for *Specificity*.

| Features |  | Acc | Sns | Spc |
| --- | --- | --- | --- | --- |
| EC | naive | 0.57 | 0.54 | 0.60 |
| FC | naive | **0.79** | 0.74 | 0.83 |
| PL | (DPH) | 0.44 | 0.39 | 0.50 |
|  | (PH) | 0.48 | 0.49 | 0.47 |
| PI | (DPH) | 0.49 | 0.44 | 0.53 |
|  | (PH) | 0.56 | 0.59 | 0.52 |
| CC | (DPH) | 0.34 | 0.18 | 0.51 |
|  | (PH) | 0.61 | 0.57 | 0.64 |
| PE | (DPH) | 0.54 | 0.67 | 0.41 |
|  | (PH) | 0.56 | 0.46 | 0.67 |

We observe that the topologically-based classifications are close to random (with accuracy around 0.5) in both the directed and undirected approach. Hence, the schizophrenia patients are not topologically discriminated from the healthy subjects, although the FC-classification is relatively successful (with accuracy 0.79). In this case, the classification task based on the EC matrix itself did not achieve good results (accuracy 0.57, close to the performances of the directed topological features).

### 4.2. Second Dataset: scalp EEG - Epilepsy

Following the same pipeline as in the previous example, we present now a classification analysis concerning a different disease (epilepsy) and a different type of data (EEG). We use for this analysis the EEG recordings described in Sec. 3.1.

Our analysis of epileptic brain states is divided in the two cases reported below: we first assess a classification by the basic approach as applied to the fMRI data, as described in Sec. 4.2.1. Subsequently we report (in Sec. 4.2.2) the results of the analysis carried on the same dataset but involving the additional cleaning steps described in Sec. 3.6.

#### 4.2.1. EEG analysis - default approach

In this analysis, we compare the functional and effective connectivity networks associated to the ictal and to the inter-ictal periods. We consider 18 subjects; each subject has both ictal and interictal data segments. For details, see Sec. 3.1.2. We apply the SVM classifier to the masked EC directed graphs, to the masked FC undirected graphs and to the topological features PL, PI, CC and PE obtained by the PH computation on both the obtained networks. We report the achieved classification values in Table 2.

**Table 2:** Epilepsy: group analysis, 18 subjects. SVM classification results using FC, EC and the topological features PL, PI, CC, PE in dimension 1 with undirected approach (PH) or directed approach (DPH). *Acc* stands for *Accuracy, Sns* for *Sensitivity* and *Spc* for *Specificity*.

| Features |  | Acc | Sns | Spc |
| --- | --- | --- | --- | --- |
| EC | naive | **0.83** | 0.72 | 0.94 |
| FC | naive | 0.78 | 0.72 | 0.83 |
| PL | (DPH) | 0.50 | 0.50 | 0.50 |
|  | (PH) | 0.50 | 0.44 | 0.56 |
| PI | (DPH) | 0.47 | 0.44 | 0.50 |
|  | (PH) | 0.64 | 0.50 | 0.78 |
| CC | (DPH) | 0.44 | 0.78 | 0.11 |
|  | (PH) | 0.69 | 0.67 | 0.72 |
| PE | (DPH) | 0.53 | 0.44 | 0.61 |
|  | (PH) | 0.33 | 0.39 | 0.28 |

The results show that the classifications based on the topological features perform again close to random, even though both the raw EC-based and raw FC-based classification are quite effective (with accuracy of 0.83 and 0.78). Therefore, when applied to both the functional and the effective connectivity networks and when compared with the results achieved by the raw matrices, we do not see any advantage in using the topological features.

However, we conjectured that the low performance of the persistent homology approach may be due to its high sensitivity to baseline inter-subject variability in the connectivity structure. Indeed, when applied to the discrimination of ictal and interictal states within a single subject, TDA provides good results, comparable in some cases to those achieved by EC or FC itself, see Appendix C. Therefore, further supported by this successful within-subject application, in the following group analysis we first attempt to correct for the inter-subject variability, before applying the classifier.

#### 4.2.2. EEG analysis: alternative approach

In order to reduce the inter-subject variability of the EEG dataset, in the following group analysis we apply further cleaning steps to the main pipeline, following the methodological details described in Sec. 3.6. In this analysis we use the same dataset as in Sec. 4.2.1. The obtained results, in dimension 1, are shown in Table 3; we refer to the Appendix, and in particular to Table D.9, for further results in dimension 0.

**Table 3:** Epilepsy: EEG group analysis (with variation) 18 subjects. SVM classification results using the topological features PL, PI, CC, PE in dimension 1, with undirected approach (PH) or directed approach (DPH). *Acc* stands for *Accuracy, Sns* for *Sensitivity* and *Spc* for *Specificity*.

| Features |  | Acc | Sns | Spc |
| --- | --- | --- | --- | --- |
| EC | naive | 0.44 | 0.56 | 0.33 |
| FC | naive | 0.89 | 0.78 | 1.00 |
| PL | (DPH) | 0.58 | 0.44 | 0.72 |
|  | (PH) | 0.75 | 0.72 | 0.78 |
| PI | (DPH) | 0.86 | 0.89 | 0.83 |
|  | (PH) | 0.83 | 0.83 | 0.83 |
| CC | (DPH) | 0.61 | 1.00 | 0.22 |
|  | (PH) | **0.92** | 0.94 | 0.89 |
| PE | (DPH) | 0.89 | 0.89 | 0.89 |
|  | (PH) | 0.06 | 0.06 | 0.06 |

We now see, in Table 3, that both DPH and PH achieve on average good performances, with around 70 percent accuracy on average. The directed and undirected method are comparable to FC, whereas EC performs close to random.

The results shown in Table 2 and in Table 3 suggest that the cleaning steps help at increasing the performance of the classification tasks from persistent homology features; on the other hand, the better performances could also be due to some specific properties of the considered data. Finally, we have applied the alternative pipeline to a different dataset, namely to epileptic seizure classification in intracranial EEG data.

### 4.3. Third Dataset: iEEG - Epilepsy

In our last analysis, we use the iEEG dataset described in Sec. 3.1.2. As in the previous section, in order to reduce the inter-subject variability, we apply the cleaning steps of Sec. 3.6 before computing the homology features and the classification. The obtained results, are shown in Table 4 and we refer to Table D.9 to additional results in dimension 0.

**Table 4:** Epilepsy: iEEG analysis, 16 subjects. SVM classification results using the topological features PL, PI, CC, PE in dimension 1, with undirected approach (PH) or directed approach (DPH). *Acc* stands for *Accuracy, Sns* for *Sensitivity* and *Spc* for *Specificity*.

| Features |  | Acc | Sns | Spc |
| --- | --- | --- | --- | --- |
| PL | (DPH) | 0.50 | 0.56 | 0.44 |
|  | (PH) | 0.62 | 0.69 | 0.56 |
| PI | (DPH) | 0.62 | 0.62 | 0.62 |
|  | (PH) | 0.66 | 0.69 | 0.62 |
| CC | (DPH) | 0.78 | 0.81 | 0.75 |
|  | (PH) | 0.84 | 0.87 | 0.81 |
| PE | (DPH) | 0.69 | 0.69 | 0.69 |
|  | (PH) | **0.87** | 0.87 | 0.87 |

The results show that good classification performances in the detection of the brain alterations have been achieved, *e.g*., with values of 0.94 for PE (in dimension 0, see Table E.11) and 0.87 obtained by PE in dimension 1. These are comparable to the results obtained in the alternative pipeline for scalp EEG data analysis of Sec. 4.2.2 and the single subject analysis presented in Appendix C and consolidate the idea that the cleaning step procedures may help in the analysis of brain state detection via TDA. out here that, in this case, a direct comparison with the performances of the connectivity matrices across subjects via FC/EC is not applicable due to interindividual differences in the electrodes number and placement.

For the sake of completeness, we report in Tab. E.11 in the appendix also other detailed results of this last analysis, such as employing the full (not-masked) matrices as well and further 0-dimensional features.

## 5. Discussion

In this work we have investigated the employment of a directed extension of persistent homology to classification tasks of disease-related alterations of brain connectivity from neuroimaging data in the case of two typical brain connectivity alterations, namely an easy task classification between ictal and pre-ictal states, and a more difficult task of classification between schizophrenia patients and healthy controls. The results of the classification of ictal from inter-ictal effective connectivity have provided a proof of principle that the directed persistent homology approach allows extracting features that carry relevant information about the underlying brain state in a real-world setting. However, the failure at the schizophrenia classification task pointed to limitations of the approach. There are several factors that play role here. First, the results of the ‘naive’ classification using directly the raw connectivity matrices suggest, that it might be that in the case of the fMRI schizoprenia classification task, the estimated effective connectivity (unlike functional connectivity) might have not contained or retained sufficient information to reliably distinguish the patients from the control. In general, effective connectivity aims for richer information concerning the system than functional connectivity (in particular the direction, and in principle the full information about the causal interactions, that can be considered the primary object that gives rise to the secondary statistical dependences captured by the functional connectivity). However, effective connectivity pays for that with higher difficulty of *estimating* it reliably. This may thus also partially explain why even in the seizure classification tasks, the PH approach typically demonstrated similar or better classification performance than the DPH approach, although among the analyses, both situations appear: PH performing better than DPH and vice versa. Indeed, it seems that the most important factor that influences the performances of the classifier is the dataset. We refer to the Appendix, Section F for a more detailed discussion of the statistical comparison of the two pipelines, containing permutation-based tests of the classification performances against the null hypothesis of random class assignment, and testing the hypotheses of the EC versus the FC-based classifiers with corresponding feature selection.

As mentioned earlier, the results generally suggest that the naive classifications based on functional or effective connectivity generally perform similarly or better than the topologically-based ones. The better performances of FC/EC on the top of TDA might be due to various reasons, including potentially a lower sensitivity of Persistent Homology to small localized variations of the absolute values of the input adjacency matrices: small variations of the metric properties of relatively small subgraphs, like the distances associated to a subset of nodes and represented by the entries of the adjacency matrix, are loosely distinguished by the associated topological features, but the same variations are better captured by classification methods based on the full raw matrices. This is amplified in the heterogeneous group-studies; in Appendix C, we report an analysis of the EEG epileptic data restricted to the single subjects showing better performances than the group-studies. Driven by the idea that TDA methods would be more effective when tested on more homogeneous data, in the second part of our analysis on the EEG datasets, we have introduced further steps in the data processing and cleaning (see Sec. 3.6). A theoretically preferable procedure would require to use the covariance matrices instead of the correlations, because satisfying linearity properties and carrying more biological information, but this would involve numerical complications. We observe that, when following this variation of the pipeline, both our computations on the EEG and iEEG data achieve slightly better performances, suggesting that the cleaning procedures may in fact be of importance in the analysis of these data.

Another observation concerns the field of applicability of the persistent homology: for some datasets (as for iEEG data), a classification analysis based on the raw FC/EC matrices can not be undertaken. This is because the analysis assumes fixed number and position of the signal sources, an assumption clearly not fulfilled by the placement of the intracranial electrodes in different subjects. Although under some circumstances homogenous resampling might be attempted (such as averaging of signals within anatomical regions), investigations like those shown on the iEEG data might be the natural field of application of TDA within neuroimaging. We note here that other measures such as raw FC/EC with some dimension reduction or indeed other methods not using connectivity at all are likely to provide similar or even higher classification performance than that achieved by the current combinations of connectivity and persistent homology features, as the seizure network dynamics are indeed commonly quite different from the inter-ictal period. We refer readers interested in this particular classification task to a recent review by Siddiqui et al. (2020), however noting that for a range of the reviewed studies, only the best performing classifier is reported, and the accuracies might therefore be generally inflated.

Due to richness of the available methodologies, we necessarily had to limit our investigation by making some specific choices. Among the general advantages of PH over more classical methods, PH does not depend upon the choice of a threshold. However, it does depend on the choice of the filtrations used and simpler graph filtrations (see, *e.g*., Lee et al. (2011a)) may also be employed as input of PH instead of the more general Rips/clique filtration. Although both the graph and Rips/clique filtrations lead to the same features in dimension 0 (as the number of connected components of the graphs and of the associated simplicial complexes is the same), this is not true in higher dimensions. In fact, the *n*-dimensional features obtained by using the graph filtration are 0 for every *n* ≥ 2, whereas, due to the presence of higher-order simplices, when using the Rips/clique filtration non-trivial *n*-dimensional holes may arise. In this work we have used the more general clique filtration, that in principle provides additional information due to the presence of higher-order simplices; however this does not necessarily mean that the classifiers will be able to use this information efficiently; however, we have not provided a comparison with other filtrations. Among other specific choices, for instance, many more featurizations of persistent homology are available in the literature, and we thus decided to focus only on some of the more common. Among other examples, kernel methods are also prominent, and we report some kernel-based classifications in Appendix D. The results reported in Section 4 and in the Appendix, refer to computations using features in dimension 0 and 1; we have also computed the persistent homology groups in higher dimensions, but, due to the size of our datasets, the Betti numbers were typically zero (and always zero from dimension 3 higher), so we did not include them in the report. It is an open question whether, for larger datasets, the higher Betti numbers can be effectively used for reaching better classifications.

Moreover, there is the choice of potentially discarding the weak (potentially noise-related) links from consideration. Among prior studies analysing functional connectivity networks through persistent homology methods, some set statistically insignificant correlations (with p-values greater than 0.05) to 0 (see Merelli et al. (2016); Stolz et al. (2018)) and others do not (see Lee et al. (2011b, 2012); Rathore et al. (2019)). The results reported in this work refer to connectivity networks with links surviving significance threshold of *α* = 0.05, but either approach can be equivalently undertaken; for sake of completeness, we report in Table D.7 the results obtained by using the full EC or FC matrix (from which we compute the distance matrix by the transformation 1 − |·|) and a comparison of the performances, showing that the performance does not substantially depend on the choice of these masks. For consistency with the undirected approach, in our study of the effective connectivity networks we introduce a similar masking approach: we first build the (pair-wise) Granger Causality graph associated to the brain data and then we set to 0 the connections in the network marked as dependent by applying the FACDA algorithm (Kořenek and Hlinka, 2020). Table D.7 shows that also in the case of the effective network, masking does not change the results substantially.

Another specific choice is the use of the SVM rather than other classifier. As mentioned in the Methods section, SVM is a relatively standard method of choice for similar data situations, used also previously in similar context Rathore et al. (2019); Stolz et al. (2018), although a plethora of alternatives exist. As with most classification tools, the performance of SVM might be adversely affected by high dimensionality of input data. However, except for PL and, potentially, PI, all the other TDA features that we have used are low-dimensional, in particular in comparison to the number of features in the naive approach that uses the full connectivity matrix as its input.

In order to study effective connectivity networks from a topological point of view, there are two approaches: one can either construct an undirected graph from the directed effective connectivity network (hence losing some information) and then apply the usual PH pipeline or one can use a suitable directed version of PH. To the best of our knowledge, this is the first attempt at employing the latter, directed PH approach to analyse effective connectivity. However, this approach has been foreseen in a recent perspective paper by Lee (2019), suggesting that causal connectivity methods are to be examined and applied for classification of diseases.

To summarise, we have shown that the (directed) persistent homology approach can be applied to brain functional/effective connectivity, obtaining in some cases good performances. However we suggest that a due caution in the use of TDA and of these featurizations should be taken, in line with very recent critical results on TDA based on functional connectivity in the literature Rathore et al. (2019); Ellis et al. (2019).

## 6. Conclusion

In this work we have shown that in principle the directed extension of persistent homology can be successfully applied to classification of disease-related alterations of brain connectivity from neuroimaging data. However, cautious direct comparison to the performance of standard (undirected) persistent homology features as well as to the use of the raw effective (or functional) connectivity matrices for the same classification task suggests that there are important challenges to be overcome to fully utilize the theoretical promise of the topological data analysis. We conclude that tackling the identified real-world neuroimaging challenges such as topographical versus topological specificity of the brain alterations, inter-individual biases and homogeneous versus heterogeneous sampling situations, would be necessary in order to firmly establish TDA also in the particular field of neuroimaging as a pragmatic tool.

## Acknowledgment

The authors thank Jakub Kopal for sharing the preprocessed fMRI time series and Barbora Bučková for sharing scripts for classification pipeline.

## A. Background: Topological Data Analysis

Topological Data Analysis is a recent and active research field of applied algebraic topology, whose main goal is to study the structure of datasets by means of topological frameworks. Among the tools of TDA maybe the most common is nowadays Persistent Homology, whose definition is based on the mathematical ideas of *simplicial complexes, filtrations* and *homology*. In this section we present a brief overview on this subject, we describe these concepts in the sections Sec. A.1 and Sec. A.2 below (referring to Zomorodian and Carlsson (2005); Otter et al. (2017) for a mathematical introduction and to Petri et al. (2013); Horak et al. (2009) for further studies of Persistent Homology with applications to complex networks).

### A.1. Simplicial complexes

Simplicial complexes are combinatorial mathematical objects that generalize (in higher dimensions) the classical definition of graphs. Roughly speaking, a simplicial complex is a space built by assembling together points, edges, triangles, tetrahedra and, more generally, high dimensional polytopes (in topology called simplices). We refer to Munkres (1984) for more details.

#### Definition A.1.

Let *V* be a set. An *(abstract) simplicial complex* on *V* is a collection Σ = {*σ*_*α*_} of finite non-empty subsets *σ*_*α*_ of *V* such that:

i. for each *v* ∈ *V*, {*v*} belongs to Σ;
ii. if *σ* ∈ Σ and *τ* ⊆ *σ*, then *τ* ∈ Σ.

The elements *σ*_*α*_ of a simplicial complex Σ are called *simplices* of Σ and if *σ*_*β*_ is a subset of a simplex *σ*_*α*_, then we say that *σ*_*β*_ is a *face* of *σ*_*α*_. The *dimension* of a simplex *σ* is one less its cardinality, i.e., if *σ* = {*v*_0_, …, *v*_*k*_}then dim *σ* = *k*, and we say that *σ* is a *k*-simplex. The 0-simplices {*v*} ∈Σ are called the *vertices* of the simplicial complex Σ. We will write *V* (Σ) for the set of vertices of Σ; with abuse of notation, we will make no distinction between the element *v* ∈ *V* and the vertex {*v*} ∈ *V* (Σ).

In this work we consider only finite simplicial complexes, so that the family Σ = {*σ*_*α*_} is finite. The dimension of the simplicial complex Σ is the maximal dimension across all its simplices.

#### Observation

Definition A.1 is rather abstract. However, one can always geometrically realize an (abstract) simplicial complex as a geometric sub-space of ℝ^*n*^. A *k*-simplex *σ* = {*p*_0_, …, *p*_*k*_} can be geometrically depicted as the convex hull of *k*+1 geometrically independent points *p*_0_, …, *p*_*k*_ in ℝ^*k*+1^. For example, a 0-simplex is a point, a 1-simplex is depicted as a segment (a convex hull of its two end-points, which are again 0-simplices). Likewise, a 2-simplex is geometrically understood as a triangle, and so on. Roughly, for geometrically depicting a simplicial complex, one realizes each of its simplex as a subspace of ℝ^*n*^ and then glues them to each other along common faces. See Fig. A.7 for an example of geometric realizations of different simplicial complexes (there denoted by Σ_0_, …, Σ_5_).

##### Example A.2.

Recall that a graph *G* is a pair *G* = (*V, E*) given by a set *V* of vertices and a set *E* ⊆*V* × *V* of edges, where an edge is identified by the couple (*v, w*) of its distinct endpoints (so we do not allow multiple edges or loops). Hence, a graph is a simplicial complex whose vertex set is *V* and the 1-simplices are the edges of *G*.

If Σ and Σ′ are simplicial complexes, a simplicial map Σ →Σ′ between them is a function *f*: *V* (Σ) → *V* (Σ′) sending vertices of Σ to vertices of Σ′, with the property that, if the vertices *v*_0_, …, *v*_*n*_ span a simplex *σ* = {*v*_0_, …, *v*_*n*_} of Σ, then also the vertices *f* (*v*_0_), …, *f* (*v*_*n*_) span a simplex of Σ′; observe that the image of *σ* can be a simplex of Σ′ of smaller dimension. Examples of simplicial maps arise from inclusion. For example, if Σ is contained in Σ′ (as subfamily) then the inclusion Σ ⊆ Σ′ defines a map of simplicial complexes (sending the simplices of Σ to the same simplices seen in Σ′. In this case we say that Σ is a *subcomplex* of Σ′. Having maps between simplicial complexes preserving the structures is an important ingredient in developing the theory of filtrations and persistent homology (see the sections below).

##### Example A.3

(Clique complex). A graph *G* is a 1-dimensional simplicial complex (as its simplices are at most of dimension 1); recall that an *n*-clique is a complete subgraph of *G* on *n* vertices. The *clique complex Ĝ* associated to *G* is the simplicial complex constructed as follows:

1. its 0- and 1-simplices are the vertices and edges of *G*;
2. for every (*n* + 1)-clique with vertices {*v*_0_, …, *v*_*n*_} in *G* we add the *n*-simplex *σ*_*α*_ = {*v*_0_, …, *v*_*n*_} on the same vertices.

For example, if *G* is a complete graph on 3 vertices, then its clique complex is nothing but a simplicial complex consisting of a 2-simplex together with all its faces. Observe that the dimension of the clique complex depends on the dimension of the maximal cliques of *G*. The embedding of *G* into *Ĝ* sending the vertices and edges of *G* to the respective 0 and 1-simplices of *Ĝ* is a map of simplicial complexes.

We are ready to recall the definition of filtration of a simplicial complex Σ:

##### Definition A.4.

A *filtration ℱ* (of a simplicial complex Σ) is an indexed sequence of subcomplexes of Σ

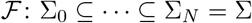

such that for every *i* ≤ *j*, the simplicial complex Σ_*i*_ is contained in Σ_*j*_.

We will only consider finite filtrations of simplicial complexes (so that the chain of simplicial complexes in the definition is finite). The set of indices in the definition of a filtration is usually given by positive natural numbers, but it can also be a more general set, like the positive real numbers (see, *e.g*., Example A.6). A figurative illustration of a filtration (with clique complexes) is given in Fig. A.7.

**Figure A.7:**
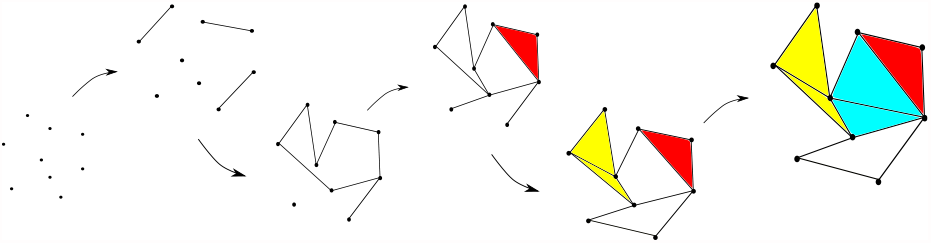
Example of a filtration of simplicial complexes. At each step we add some edges, and when we find a 3-clique we fill it in with a (coloured) 2-simplex.

We conclude the subsection with two prominent examples of filtrations: the *weight rank clique filtration* (WRCF) Petri et al. (2013) associated to a weighted graph and the *Vietoris–Rips filtration* associated to any finite subset of a metric space.

##### Example A.5

(WRCF). Let *G* be a weighted graph and let (*w*_*i*_)_0≤*i*≤*N*_ be the increasing sequence of all the weights of *G* (i.e., *w*_0_ is the minimal weight, *w*_*N*_ the maximal and *w*_*i*_ *< w*_*i*+1_). For the *i*-th weight *w*_*i*_ construct the simplicial complex Σ_*i*_ as follows:

1. consider the subgraph *G*_*i*_ ⊆ *G* with the same vertices of *G* and only those edges of *G* with corresponding weight *w* such that *w* ≤ *w*_*i*_;
2. Σ_*i*_ := *Ĝ*_*i*_ is the clique complex of *G*_*i*_ (see Ex. A.3).

It is easy to see that Σ_*i*_ is contained in Σ_*i*+1_ and that the sequence Ø ⊆ Σ_0_ ⊆ · · · ⊆ Σ_*N*_ = Σ is a filtration. Analogously, by sorting the weights in the decreasing order, one obtains another filtration, where the graph *G*_*i*_ is defined by keeping only the edges of *G* with weight *w* ≥ *w*_*i*_.

##### Example A.6

(Vietoris–Rips filtration). Let (*X, d*) be a metric space and *S* be a finite subset of *X*, endowed with the induced metric. In applications, the set *S* can be a set of data-points in ℝ^*n*^ or a set of measurements with a notion of distances between them. The *Vietoris–Rips complex of S at scale t* ≥ 0 is the simplicial complex:

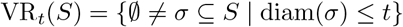

where the diam(*σ*) := sup{*d*(*x, y*) |*x, y*∈ *σ*} is the diameter of *σ* (we point out here that different definitions, considering a maximal distance of 2*t* instead of *t* can be found in the literature; we follow the definition used in Tralie et al. (2018) to be consistent with Sec. 3.8). Observe that every such *σ* represents a simplex of dimension |*σ*| − 1. If *t* ≤ *t*^/^ then VR_*t*_(*S*) ⊆ VR(*S*)_*t′*_ and the resulting filtration VR(*S*) of the simplicial complexes VR_*t*_(*S*), with *t* varying in ℝ_≥0_, is called the *Vietoris–Rips filtration* of *S*.

##### Remark A.7.

Observe that the two constructions in Ex. A.5 and in Ex. A.6 are equivalent, if one considers metric spaces (*X, d*) associated to weighted graphs *G* (with respect to an increasing ordering of the weights), with underlying space *X* given by the nodes of the graph *G* and distance *d*(*x, y*) between *x* and *y* defined as the weight corresponding to the edge (*x, y*).

### A.2. (Persistent) Homology and Persistence diagrams

Persistent Homology (PH) Edelsbrunner et al. (2002); Zomorodian and Carlsson (2005) is one of the main tools in TDA and one of the first applications of Algebraic Topology to Data Analysis, introduced in Carlsson (2009). Persistent Homology is a topological invariant whose definition is based on the mathematical concept of homology groups.

It is robust to noise and stable with respect to small perturbations of the inputs (Cohen-Steiner et al., 2007, Section 3.1); moreover, calculations can be easily achieved by using many software options (in MATLAB, Python, C++, R, Java) Otter et al. (2017). Its constituents, the homology groups, are classical algebraic invariants of topological spaces. The number of connected components, loops (1-dimensional holes), voids (2-dimensional holes), tunnels and higher dimensional holes are examples of such invariants. A technical comprehensive definition of homology groups is outside of the scope of the present work. In the following, we will only sketch some ideas, but the interested reader can find more details in Hatcher (2000); Munkres (1984). We conclude the section with the definition of persistent homology groups and persistence diagrams.

Let *k* be a natural number. The *k*-th *homology group H*_*k*_(Σ) associated to a simplicial complex Σ (see Def. A.1) can be thought of as the set (group, module or vector space) of *k*-dimensional holes in Σ; the elements of *H*_∗_(Σ) are called *homology classes*. To be more precise, let 𝔽_2_ be the field with two elements (i.e., 0 and 1, with sum and product inherited from the usual sum and product of real numbers, reduced mod 2) and let *C*_*p*_(Σ) be the free 𝔽_2_-vector space whose basis consists of the set of *p*-simplices of Σ: elements of *C*_*p*_(Σ) are formal combinations *a*_0_*σ*_0_ + · · ·+*a*_*n*_*σ*_*n*_ of *p*-simplices *σ*_0_, …, *σ*_*n*_ of Σ with coefficients *a*_*i*_ in 𝔽_2_. For every *p* ≥ 1 we define the map

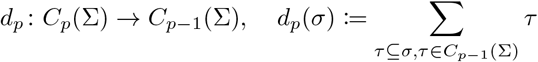

by sending a *p*-simplex *σ* to a formal sum of all its (*p* − 1)-faces. The map *d*_0_ is defined as the zero map. As the composition *d*_*p*_ ∘ *d*_*p*+1_ = 0 is the zero map, the image of *d*_*p*+1_ in *C*_*p*_(Σ) is contained in the kernel of *d*_*p*_; these are both 𝔽_2_-vector spaces and the quotient ker(*d*_*k*_)*/*Im(*d*_*k*+1_) of vector spaces is well-defined. We can now formally define the homology groups of the simplicial complex Σ (with 𝔽_2_-coefficients):

#### Definition A.8.

Let Σ be a simplicial complex and *k* ≥ 0 a natural number. The *k*-th *homology group H*_*k*_(Σ) of Σ

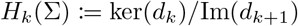

is defined as the quotient of the kernel of the map *d*_*k*_ with the image of *d*_*k*+1_.

The group *H*_*k*_(Σ) is in fact a 𝔽_2_-vector space and its dimension (over 𝔽_2_) called the *k*-th *Betti number β*_*k*_(Σ) of Σ. More general definitions are possible, with arbitrary fields 𝔽 or with the integers ℤ as coefficients. We refer to Hatcher (2000); Munkres (1984) for more detailed descriptions of homology groups.

The number of *k*-holes of (the geometric realization of) a simplicial complex Σ corresponds to the *k*-th Betti number *β*_*k*_(Σ) associated this way to Σ. In fact, Betti numbers give (a measure of) the number of “independent” holes or loops in Σ, i.e., holes that can not be shrunk continuously to a point. The 0-th Betti number *β*_0_ represents the number of connected components, the 1-st Betti number *β*_1_ is the number of independent loops and the 2-nd Betti number *β*_2_ gives the number of 2-dimensional voids.

#### Example A.9.

If *X* is a point, then *X* has a single connected component and no loops or higher dimensional holes. Its Betti numbers *β*_*n*_(*X*) are 0 for every *n >* 0, except for *β*_0_(*X*) which is 1. The 0-th Betti number of two points is 2 (2 connected components) and 0 in higher dimensions. On the other hand, if *X* is a 2-dimensional sphere, then *X* has a 2-dimensional hole, represented by the void internal to the sphere. We get *β*_0_(*X*) = *β*_2_(*X*) = 1 and *β*_*n*_(*X*) = 0 otherwise. Observe that *β*_1_(*X*) is also 0 as every loop on the sphere can be continuously shrunk to one point.

One of the most useful theoretical properties of homology is given by its functoriality: for every simplicial map *f*: *X* → *Y* of simplicial complexes and for every *k* in ℕ, we also get a linear map *H*_*k*_(*f*): *H*_*k*_(*X*) → *H*_*k*_(*Y*):

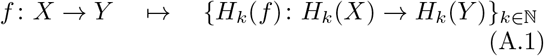

satisfying some additional compatibility properties; we avoid to spell them out here and we refer to Hatcher (2000) for the details. This turns out to be of great importance when dealing with more than one single simplicial complex.

As we have seen, the *k*-th homology group *H*_*k*_(Σ) is associated to a single simplicial complex Σ. Likewise, persistent homology groups are defined for a filtration of simplicial complexes, as we now explain. By the functoriality described in Eq. A.1, if ℱ: Ø⊆Σ_0_ ⊆· · ·⊆Σ_*N*_ = Σ is a filtration of simplicial complexes (see Def. A.4), then, for every *k*, we get a sequence 0 → *H*_*k*_(Σ_0_) → · · · → *H*_*k*_(Σ_*N*_) = *H*_*k*_(Σ) of homology groups and morphisms between them. Let *k* be fixed; then, for every pair of indices *i* and *j* between 0 and *N*, the composition of maps of simplicial complexes Σ_*i*_ → · · · → Σ_*j*_ induces a map 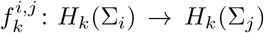 between the homology groups of Σ_*i*_ and Σ_*j*_.

#### Definition A.10.

Edelsbrunner et al. (2002) For *i < j*, the *k*-th *persistent homology group* 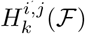 of the filtration ℱ: Ø ⊆ Σ_0_ ⊆· · · ⊆ Σ_*N*_ = Σ is the image of the induced map 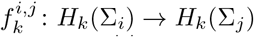. The *k*-th *persistent Betti numbers* 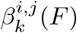 are the ranks of the persistent homology groups 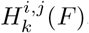.

The intuition behind the definition of persistent homology groups is that homology classes (the *k*-dimensional holes) can appear and disappear at different stages of the filtration steps, due to changes in the topology of the simplicial complexes across the filtration. A persistence homology class (i.e., an element) of 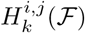 represents a *k*-dimensional hole of Σ_*i*_ surviving across all the filtration steps between the index *i* and index *j*. The filtration indices at which a *k*-dimensional hole first appears and first disappears are called its *birth* and *death*. Persistent homology makes (mathematically) clear the idea of appearing and disappearing of high dimensional holes; it keeps track of all these topological changes with the persistence Betti numbers.

We can visualize the information provided by persistent homology groups and persistent Betti numbers by constructing a planar diagram called *Persistence Diagram* (PD), stably with respect to the input data Cohen-Steiner et al. (2007). A persistent homology class born at time *b* and that died at time *d* is represented in the diagram by the pair (*b, d*) (see Fig. A.8). Classes that persist until the end are usually depicted in the persistent diagram as dying at infinity (or at any other index bigger than the maximum filtration one). Note that there are no points below the diagonal because the death value of a hole is always bigger than its birth value.

In a persistence diagram it is then visually clear the measure of how long a homology class persists: the farther from the diagonal, the higher the persistence. In the standard interpretations, short-living classes (so, with representing dots close to the diagonal) are interpreted as noise, and only long-persistence classes represent important features of the filtration.

**Figure A.8:**
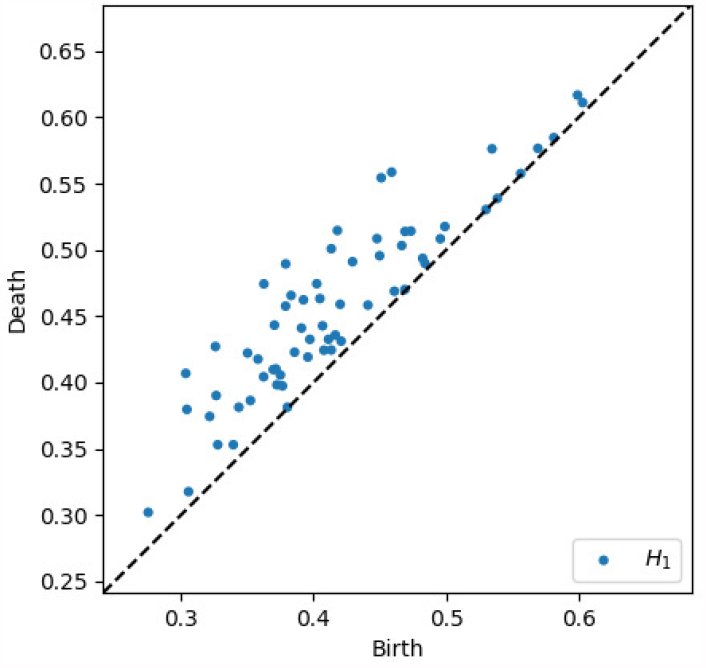
Persistence diagram associated to a filtration of simplicial complexes. Each dot represents the birth-death of a *k*-dimensional hole. The bottom right triangular segment is empty because the death of a persistent homology class can not be lower than its birth.

## B. EEG data: Subject selection

In our analysis of epileptic brain states we have used the data available at https://physionet.org/content/chbmit/1.0.0/, from the publicly open dataset *Physionet*. We have analysed EEG recordings belonging to 18 selected subjects: chb01-05, chb07, chb09, chb10, chb12, chb14, chb17-24 except chb18_01, chb19_01.

For the variation of the main pipeline, we have used the following files: chb01_03, chb02_16+, chb03_01, chb04_05, chb05_06, chb07_12, chb09_06, chb10_12, chb12_06, chb14_11, chb17a_03, chb18_29, chb19_28, chb20_13, chb21_19, chb22_20, chb23_06, chb24_04. For those, we always use the first seizure (the first 30 seconds of it) and two 30-second segments preceding it: [-3m:-2m30sec] for baseline connectivity, and [-1min:-30sec] for preictal connectivity estimation. Detailed procedure is described in Sec. 3.1.2, in Intracranial EEG data description.

## C. EEG data: single subject analysis

In this supplementary, we assess the efficiency of PH and DPH on single subjects brain state alteration classification, in contrast to the group analysis reported in the main paper in Sec. 4.2.1.

In the single-subject analysis, we consider the data collected from the patients CHB06, CHB15 and CHB24 (from the same database, we have chosen the patients with higher number of seizures).

For every patient, we consider segments with epileptic seizures and interictal data of corresponding length: 10 ictal plus 10 interictal recordings for the patient CHB06, 20 plus 20 for the patient CHB15 and 16 plus 10 for the patient CHB24. We then apply the same pipeline as in the group analysis.

From the accuracy results, shown in Table C.5, we observe that the classification performances associated to single patients, based on topological features out of the directed and undirected approach, achieve better results (on average about 0.75, 0.60 and 0.70 for CHB06, CHB15 and CHB24, respectively) than in the inter-subject analysis.

**Table C.5:**
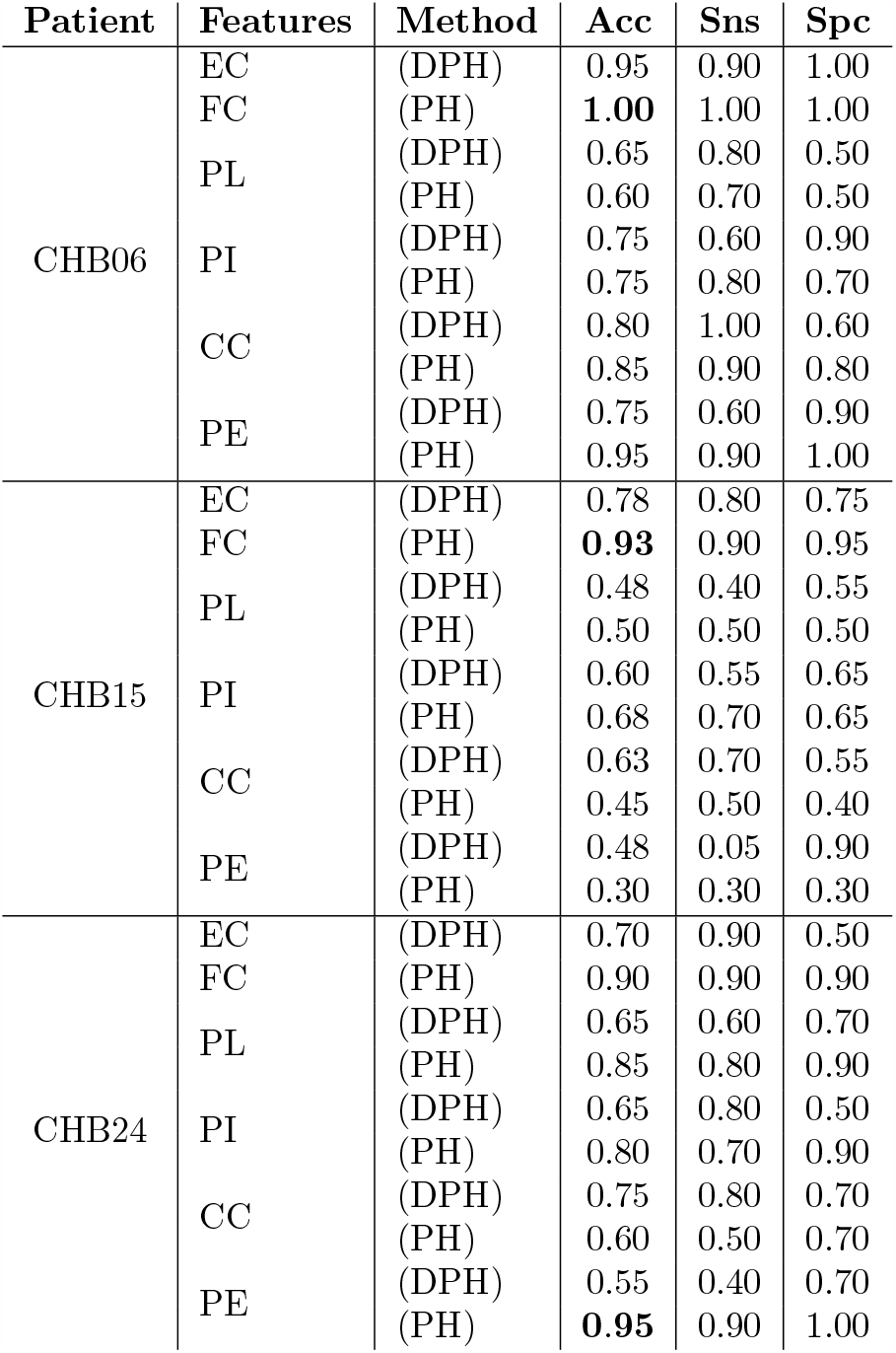
Epilepsy: single subject analysis. SVM classification of ictal and interictal data corresponding to 3 patients: CHB06, CHB15, CHB24. For each patient, the classification uses masked FC, masked EC or the topological features PL, PI, CC and PE with undirected approach (PH) and with directed approach (DPH). *Acc* stands for *Accuracy, Sns* for *Sensitivity* and *Spc* for *Specificity*.

Topological features may not achieve great performances in heterogenous group-wise classification settings (compare the results with those in Table 2), but they are more informative when applied to more specific datasets.

## D. Supplementary results

In a parallel exploratory analysis to that presented in the main text, we have applied small variations of the pipeline described in Sec. 3 and/or computed different features. In this supplementary we show the results obtained following the main pipeline applied on the fMRI and EEG datasets, using either kernel methods (see App. D.1), the full matrices (as apposed to the masked ones, see App. D.2, or the 0-dimensional features (as opposed to the 1-dimensional features, see App. D.3).

### D.1. Kernel methods

Our main analysis has been based on Persistence Images, Persistence Landscapes, Persistence Images, Carlsson Coordinates and Persistent Entropy, but many other topological features to analyse datasets are available. Among others, kernel-based methods provide some flexible choices. As there is not a standard persistence kernel approach, we have decided to analyse this feature separately. We follow the kernel-based method introduced by Reininghaus et al. (2015) and apply the SVM classification on the (masked) FC and EC networks, following the pipeline in Sec. 3. We report the results in Table D.6, from which we can see that they are comparable to those achieved by the other features reported in the main text of the work.

### D.2. Full versus masked connectivity networks

An additional analysis has been driven by the question whether classification tasks, based on masked matrices associated to (un-)directed networks (see Sec. 3.2), achieve different results when applied to the full matrices. We compare in Table D.7 full and masked-based classification results; the results suggest that the masking does not have a systematic detrimental effect on the analysis outcomes.

### D.3. 0-Dimensional features

In this analysis carried out on both EEG and fMRI data, we focus on 0-dimensional topological features. We use the features Carlsson Coordinates (CC), Persistent Entropy (PE) and the kernel feature of Reininghaus et al. (2015). Additionally, we investigate the 0-dimensional features *Area Under the Curve* (AUC) Gracia-Tabuenca et al. (2019) and *Integrated Persistence Feature* (IPF) Kuang et al. (2019a), as both have shown to provide good differentiations when applied to attention-deficit/hyperactivity disorder (ADHD) and Alzheimer’s disease, respectively. We show in Table D.8 the results on the fMRI (schizophrenia) dataset and on the scalp EEG (epilepsy) data; we present in Table D.9 the results in dimension 0 for both the scalp EEG and intracranial iEEG datasets, when using the alternative approach discussed in Sec. 4.2.2 and Sec. 4.3.

**Table D.6:** SVM kernel-based classification of functional and effective connectivity networks (both masked) using undirected approach (PH) or directed approach (DPH) (in dimension 1), respectively. Epil denotes the EEG epilepsy data for the group analysis, CHB06, CHB15 and CHB24 denote single epileptic patients and schizo denotes the fMRI schizophrenia data. *Acc* stands for *Accuracy, Sns* for *Sensitivity* and *Spc* for *Specificity*.

| Data | Method | Acc | Sns | Spc |
| --- | --- | --- | --- | --- |
| Schizo | EC (DPH) | 0.51 | 0.57 | 0.46 |
|  | FC (PH) | 0.62 | 0.67 | 0.57 |
| Epil | EC (DPH) | 0.53 | 0.39 | 0.67 |
|  | FC (PH) | 0.61 | 0.67 | 0.56 |
| CHB06 | EC (DPH) | 0.80 | 0.90 | 0.70 |
|  | FC (PH) | 0.85 | 0.80 | 0.90 |
| CHB15 | EC (DPH) | 0.43 | 0.50 | 0.35 |
|  | FC (PH) | 0.58 | 0.70 | 0.45 |
| CHB24 | EC (DPH) | 0.55 | 0.60 | 0.50 |
|  | FC (PH) | 0.75 | 0.70 | 0.80 |

**Table D.7:** Full FC vs masked FC; full EC vs masked EC. Epil denotes the EEG epilepsy data for the group analysis, CHB06, CHB15 and CHB24 denote single epileptic patients and schizo denotes the fMRI schizophrenia data. The FC/EC masks are discussed in Sec. 3.2. Features refer to the topological features of Sec. 3.5; naive denotes the classification method applied to the original matrices.

| Data | Feature | Full-FC | Masked-FC | Full-EC | Masked-EC |
| --- | --- | --- | --- | --- | --- |
| Schizo | CC | 0.59 | 0.61 | **0.59** | 0.34 |
|  | PE | 0.58 | 0.56 | 0.55 | 0.54 |
|  | PI | 0.52 | 0.56 | 0.48 | 0.49 |
|  | PL | 0.53 | 0.48 | **0.59** | 0.44 |
|  | naive | **0.80** | **0.79** | 0.54 | **0.57** |
| Epil | CC | 0.56 | 0.69 | 0.58 | 0.44 |
|  | PE | 0.56 | 0.33 | 0.61 | 0.53 |
|  | PI | 0.58 | 0.64 | 0.33 | 0.47 |
|  | PL | 0.36 | 0.50 | 0.31 | 0.50 |
|  | naive | **0.81** | **0.78** | **0.89** | **0.83** |
| CHB06 | CC | **1.00** | 0.85 | 0.70 | 0.80 |
|  | PE | 0.80 | 0.95 | 0.60 | 0.75 |
|  | PI | **1.00** | 0.75 | 0.65 | 0.75 |
|  | PL | 0.95 | 0.60 | 0.60 | 0.65 |
|  | naive | **1.00** | **1.00** | **1.00** | **0.95** |
| CHB15 | CC | 0.28 | 0.45 | 0.58 | 0.63 |
|  | PE | 0.48 | 0.30 | 0.48 | 0.48 |
|  | PI | 0.50 | 0.68 | 0.63 | 0.60 |
|  | PL | 0.73 | 0.50 | 0.65 | 0.48 |
|  | naive | **0.93** | **0.93** | **0.95** | **0.78** |
| CHB24 | CC | 0.90 | 0.60 | 0.65 | **0.75** |
|  | PE | **1.00** | **0.95** | 0.60 | 0.55 |
|  | PI | 0.75 | 0.80 | 0.40 | 0.65 |
|  | PL | 0.95 | 0.85 | 0.70 | 0.65 |
|  | naive | 0.80 | 0.90 | **0.85** | 0.70 |

## E. Variation on EEG and iEEG dataset: the complete report

We report here the full analysis based on the variation of the pipeline (as described in Sec. 3.6) in the case of the EEG/iEEG recordings. In the results of Sec. 4.2.2 and Sec. 4.3, we have only shown the classification results based on the masked FC and on the masked EC matrices (see Sec. 3.2). For the sake of completeness, we show below in Tab. E.10 and Tab. E.11 the results achieved by the topological features PL, PI, PE, CC (see Sec. 3.5), kernel (introduced in App. D.1), and by the features AUC and IPF (as in the previous section), in both dimension 0 and dimension 1 and for Full-FC, Masked-FC, Full-EC and Masked EC.

## F. Comparison of PH and DPH classification

A direct comparison between the PH and DPH performances is not straightforward given the number and intricate depeendence of the classification tasks. In Figure F.9, we present a scatter-plot where each point represents a classification of one of the datasets using one particular TDA feature (vector). The X-coordinate shows the classification accuracy obtained with DPH (computed on EC), and the Y-coordinate shows the corresponding PH (computed on FC) results. Different datasets are marked with different colours.

We observe that, in some cases, PH and DPH are comparable - shown by the points laying close to the diagonal; in other cases, PH seems to work better - upper triangle points. The cloud around (0.5, 0.5) corresponds to results close to random for both input matrices, mainly for the schizophrenia and group epilepsy studies.

To statistically evaluate the two approaches, we present in Figure F.10 the results of permutation tests. Firstly, we assess the classification significance of both pipelines, PH- and DPH-based, by permuting the obtained labels of data points. By repeating this 500 times (500 permutations) we get a distribution of random classifications; we use accuracy as the test statistic, to compute the p-value for the null hypothesis that the accuracy of our classifier is not distinguishable from random. Corresponding (uncorrected) p-values can be found in Figure F.10, top subpanel.

**Table D.8:** SVM classification with 0-dimensional topological features, for the masked FC and EC. Epil denotes the EEG epilepsy data for the group analysis, CHB06, CHB15 and CHB24 denote single epileptic patients and schizo denotes the fMRI schizophrenia data. Features (CC) and (PE) refer to the topological features of Sec. 3.5, (kernel), (AUC) and (IPF) are additional features discussed in the text.

| Data | Features | Method | Acc | Sns | Spc |
| --- | --- | --- | --- | --- | --- |
| Schizo | CC | EC (DPH) | 0.52 | 0.30 | 0.73 |
|  |  | FC (PH) | 0.49 | 0.51 | 0.47 |
|  | PE | EC (DPH) | 0.49 | 0.19 | 0.80 |
|  |  | FC (PH) | 0.49 | 0.62 | 0.37 |
|  | kernel | EC (DPH) | 0.53 | 0.30 | 0.77 |
|  |  | FC (PH) | 0.53 | 0.59 | 0.47 |
|  | AUC | EC (DPH) | 0.49 | 0.70 | 0.28 |
|  |  | FC (PH) | 0.55 | 0.56 | 0.54 |
|  | IPF | EC (DPH) | 0.45 | 0.62 | 0.28 |
|  |  | FC (PH) | 0.38 | 0.19 | 0.57 |
| Epil | CC | EC (DPH) | 0.47 | 0.50 | 0.44 |
|  |  | FC (PH) | 0.53 | 0.61 | 0.44 |
|  | PE | EC (DPH) | 0.33 | 0.00 | 0.67 |
|  |  | FC (PH) | 0.53 | 0.72 | 0.33 |
|  | kernel | EC (DPH) | 0.42 | 0.50 | 0.33 |
|  |  | FC (PH) | 0.53 | 0.67 | 0.49 |
|  | AUC | EC (DPH) | 0.44 | 0.50 | 0.39 |
|  |  | FC (PH) | 0.56 | 0.61 | 0.50 |
|  | IPF | EC (DPH) | 0.53 | 0.78 | 0.28 |
|  |  | FC (PH) | 0.56 | 0.50 | 0.51 |
| CHB06 | CC | EC (DPH) | 0.75 | 0.70 | 0.80 |
|  |  | FC (PH) | 1.00 | 1.00 | 1.00 |
|  | PE | EC (DPH) | 0.70 | 0.80 | 0.60 |
|  |  | FC (PH) | 0.95 | 1.00 | 0.90 |
|  | kernel | EC (DPH) | 0.75 | 0.80 | 0.70 |
|  |  | FC (PH) | 1.00 | 1.00 | 1.00 |
|  | AUC | EC (DPH) | 0.65 | 0.70 | 0.60 |
|  |  | FC (PH) | 1.00 | 1.00 | 1.00 |
|  | IPF | EC (DPH) | 0.80 | 0.90 | 0.70 |
|  |  | FC (PH) | 1.00 | 1.00 | 1.00 |
| CHB15 | CC | EC (DPH) | 0.30 | 0.20 | 0.40 |
|  |  | FC (PH) | 0.58 | 0.55 | 0.60 |
|  | PE | EC (DPH) | 0.60 | 0.55 | 0.65 |
|  |  | FC (PH) | 0.58 | 0.50 | 0.65 |
|  | kernel | EC (DPH) | 0.50 | 0.40 | 0.60 |
|  |  | FC (PH) | 0.38 | 0.55 | 0.20 |
|  | AUC | EC (DPH) | 0.58 | 0.45 | 0.70 |
|  |  | FC (PH) | 0.58 | 0.75 | 0.40 |
|  | IPF | EC (DPH) | 0.63 | 0.55 | 0.70 |
|  |  | FC (PH) | 0.50 | 0.20 | 0.80 |
| CHB24 | CC | EC (DPH) | 0.70 | 0.70 | 0.70 |
|  |  | FC (PH) | 0.95 | 1.00 | 0.90 |
|  | PE | EC (DPH) | 0.60 | 0.60 | 0.60 |
|  |  | FC (PH) | 1.00 | 1.00 | 1.00 |
|  | kernel | EC (DPH) | 0.75 | 0.90 | 0.60 |
|  |  | FC (PH) | 1.00 | 1.00 | 1.00 |
|  | AUC | EC (DPH) | 0.50 | 0.60 | 0.40 |
|  |  | FC (PH) | 1.00 | 1.00 | 1.00 |
|  | IPF | EC (DPH) | 0.50 | 0.60 | 0.40 |
|  |  | FC (PH) | 1.00 | 1.00 | 1.00 |

**Table D.9:**
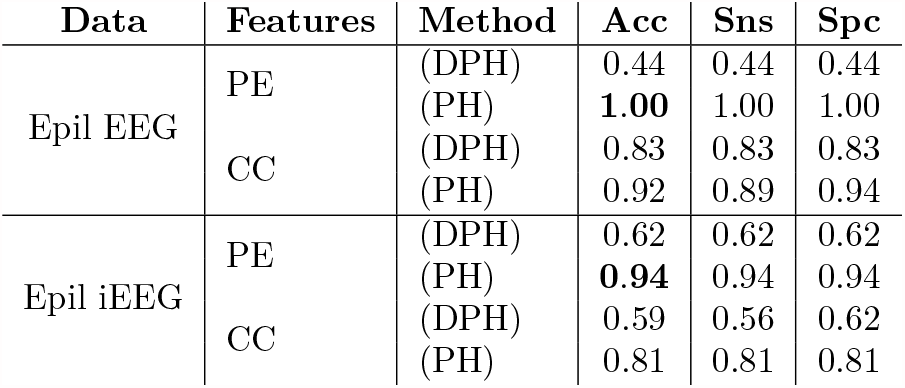
SVM classification with 0-dimensional topological features, for the masked FC and EC. Epil EEG and Epil iEEG denote the EEG epilepsy data for the group analysis for the scalp and intracranial data, respectively. Features (CC) and (PE) refer to the topological features of Sec. 3.5, (kernel), (AUC) and (IPF) are additional features discussed in the text. Reported the undirected approach (PH) or directed approach (DPH). *Acc* stands for *Accuracy, Sns* for *Sensitivity* and *Spc* for *Specificity*.

| Data | Features | Method | Acc | Sns | Spc |
| --- | --- | --- | --- | --- | --- |
| Epil EEG | PE | (DPH) | 0.44 | 0.44 | 0.44 |
|  |  | (PH) | **1.00** | 1.00 | 1.00 |
|  | CC | (DPH) | 0.83 | 0.83 | 0.83 |
|  |  | (PH) | 0.92 | 0.89 | 0.94 |
| Epil iEEG | PE | (DPH) | 0.62 | 0.62 | 0.62 |
|  |  | (PH) | **0.94** | 0.94 | 0.94 |
|  | CC | (DPH) | 0.59 | 0.56 | 0.62 |
|  |  | (PH) | 0.81 | 0.81 | 0.81 |

**Table E.10:** Epilepsy: EEG analysis with variation, full report. SVM classification with 0-dimensional and 1-dimensional topological features, for the full/masked FC and full/masked EC derived features associated to the EEG dataset, following the variation of the pipeline. The features (CC) and (PE) refer to the topological features of Sec. 3.5, (kernel), (AUC) and (IPF) are additional features discussed in the appendix.

|  | Feature | Full-FC | Masked-FC | Full-EC | Masked-EC |
| --- | --- | --- | --- | --- | --- |
| Dim 0 | AUC | 0.94 | **1.00** | 0.61 | 0.67 |
|  | IPF | 0.89 | **1.00** | 0.67 | 0.67 |
|  | PE | 0.89 | **1.00** | 0.61 | 0.44 |
|  | CC | 0.94 | 0.92 | 0.31 | 0.83 |
|  | kernel | 0.92 | 0.92 | 0.50 | 0.53 |
| Dim 1 | PI | 0.86 | 0.83 | 0.58 | 0.86 |
|  | PE | 0.33 | 0.06 | 0.50 | 0.89 |
|  | PL | 0.83 | 0.75 | 0.56 | 0.58 |
|  | CC | **0.86** | 0.92 | 0.75 | 0.61 |
|  | kernel | 0.83 | 0.81 | 0.69 | 0.86 |

**Table E.11:** Epilepsy: iEEG analysis, full report, following the variation of the pipeline. SVM classification with 0-dimensional and 1-dimensional topological features, for the full/masked FC and full/masked EC derived features. The features (CC) and (PE) refer to the topological features of Sec. 3.5, (kernel), (AUC) and (IPF) are additional features discussed in the appendix.

|  | Feature | Full-FC | Masked-FC | Full-EC | Masked-EC |
| --- | --- | --- | --- | --- | --- |
| Dim 0 | AUC | **1.00** | 0.87 | 0.62 | 0.12 |
|  | IPF | **1.00** | 0.94 | 0.75 | 0.69 |
|  | PE | **1.00** | 0.94 | 0.75 | 0.62 |
|  | CC | 0.87 | 0.81 | 0.25 | 0.59 |
|  | kernel | 0.87 | 0.87 | 0.53 | 0.50 |
| Dim 1 | PI | 0.81 | 0.66 | 0.59 | 0.62 |
|  | PE | 0.37 | **0.87** | 0.50 | 0.69 |
|  | PL | 0.78 | 0.62 | 0.47 | 0.50 |
|  | CC | **0.87** | 0.84 | 0.62 | 0.78 |
|  | kernel | 0.75 | 0.50 | 0.62 | 0.56 |

**Figure F.9:**
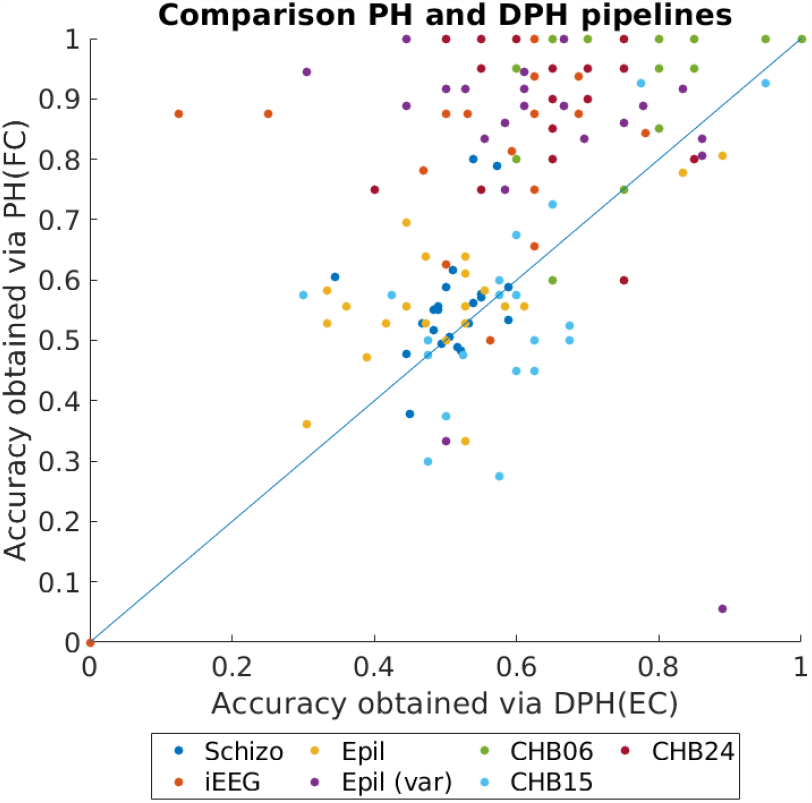
Scatter plot of classification accuracy reached using PH and DPH pipelines.

For the comparison between the DPH- and PH-based classifications, we performed another permutation test. Now, for every tested dataset and TDA feature, we work with pairs of vectors: labels for the classifier that we get from PH and from DPH. For every data point, we either flip the corresponding coordinate between these vectors, or leave it with equal probability. In this way, we generate surrogates representing the assumption “there is no difference between the PH and DPH approach performance”. We test the mentioned null hypothesis against two alternatives: PH is better than DPH, and DPH is better than PH. For that we use test statistic given by the accuracy of PH minus the accuracy of DPH, concentrating on its high positive and negative values for testing the mentioned two alternative hypotheses, respectively. Corresponding (uncorrected) p-values can be found in Figure F.10, bottom subpanel.

From the Figure F.10 it seems that the most important parameter, influencing classification, is the dataset. For our datasets, schizophrenia, epilepsy on a group level and patient CHB15 are those where the TDA accuracies are most random (not the same for the naive approaches). In the other cases, both PH and DPH seem to work better than random. We present uncorrected p-values since due to the complexity of dependencies it is difficult to choose an appropriate multiple testing correction. Both the TDA features and datasets are not independent, so it is difficult to compute the degrees of freedom for the correction. However, the number of significant classifications for both PH and DPH is clearly substantially higher than the expected 5% of false positives; so, generally, we can conclude that both pipelines work in favorable conditions. Furthermore, testing the “PH is better than DPH” alternative, seems to provide a larger number of significant classifications, which would suggest that generally PH performs better than DPH.

**Figure F.10:**
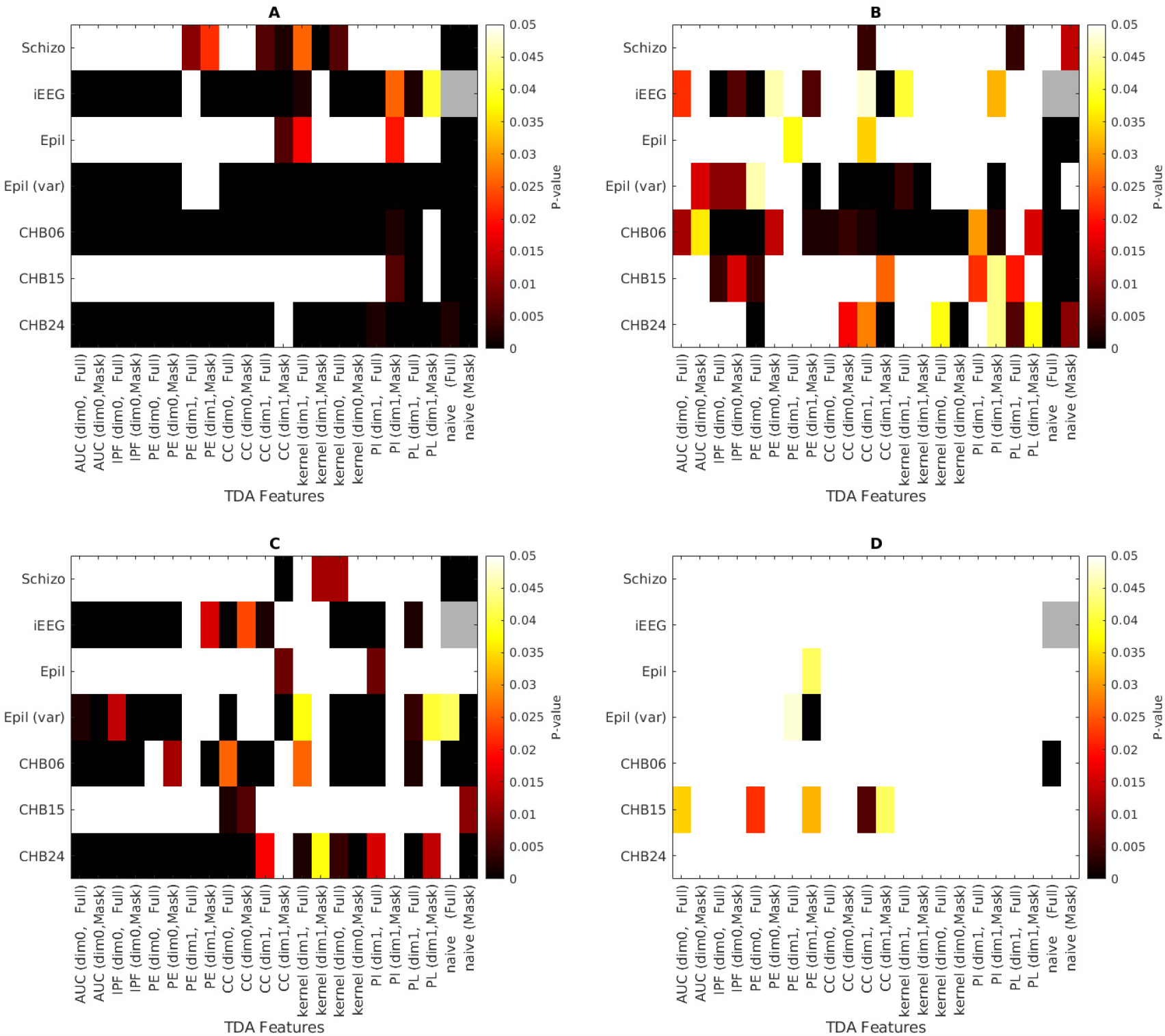
A. Uncorrected *p*-values for significance of classifications using FC(PH) pipeline. Null hypothesis: FC(PH)-based accuracy is random. Alternative: FC(PH)-based accuracy is higher than random. B. Uncorrected *p*-values for significance of classifications using EC(DPH) pipeline. Null hypothesis: EC(DPH)-based accuracy is random. Alternative: EC(DPH)-based accuracy is higher than random. C. Uncorrected *p*-values for comparison of classifications using FC(PH) and EC(DPH) pipeline. Null hypothesis: accuracy of classifiers are the same. Alternative hypothesis: FC(PH)-based accuracy is higher. D. Uncorrected *p*-values for comparison of classifications using FC(PH) and EC(DPH) pipeline. Null hypothesis: accuracy of classifiers are the same. Alternative hypothesis: EC(DPH)-based accuracy is higher. Naive classification for the iEEG dataset is not possible due to different matrix sizes across subjects.

## Notes

☆ The study was supported by the Czech Science Foundation projects No. GA19-11753S, GA21-14727K, GA21-17211S, GA21-32608S, by the Czech Health Research Council projects No. NV17-28427A and NU21-08-00432, and by Ministry of Health of the Czech Republic – DRO 2021 (National Institute of Mental Health – NIMH, IN: 00023752 “).

### Competing Interest Statement

The authors have declared no competing interest.

## References

H. Adams, T. Emerson, M. Kirby, R. Neville, C. Peterson, P. Shipman, S. Chepushtanova, E. Hanson, F. Motta, and L. Ziegelmeier. Persistence images: A stable vector representation of persistent homology. Journal of Machine Learning Research, 18(8):1–35, 2017.

Aaron Adcock, Erik Carlsson, and Gunnar Carlsson. The ring of algebraic functions on persistence bar codes. Homology, Homotopy and Applications, 18, 04 2013. doi: 10.4310/HHA.2016.v18.n1.a21.

Mehmet E. Aktas, Esra Akbas, and Ahmed El Fatmaoui. Persistence homology of networks: methods and applications. Applied Network Science, 4(61), 2019. doi: 10.1007/s41109-019-0179-3.

L. Barnett and A. Seth. The MVGC Multivariate Granger Causality Toolbox: A New Approach to Granger-causal Inference. Journal of neuroscience methods, 223, 11 2013. doi: 10.1016/j.jneumeth.2013.10.018.

P. Bendich, J. Marron, E. Miller, A. Pieloch, and S. Skwerer. Persistent homology analysis of brain artery trees. The Annals of Applied Statistics, 10, 11 2014. doi: 10.1214/15-AOAS886.

Jacek Brodzki, Francisco Belchí, Ratko Djukanovic, Joy Conway, Mariam Pirashvili, and Michael Bennett. Lung topology characteristics in patients with chronic obstructive pulmonary disease. Scientific Reports, 8, 03 2018. doi: 10.1038/s41598-018-23424-0.

Peter Bubenik and Pawe-l D-lotko. A persistence landscapes toolbox for topological statistics. Journal of Symbolic Computation, 78:91 – 114, 2017. ISSN 0747-7171. doi: https://doi.org/10.1016/j.jsc.2016.03.009. Algorithms and Software for Computational Topology.

Ed Bullmore and Olaf Sporns. Complex brain networks: graph theoretical analysis of structural and functional systems. Nature Reviews Neuroscience, 10(3):186–198, 2009. doi: 10.1038/nrn2575.

G. Carlsson. Topology and data. Bull. Amer. Math. Soc., 46:255–308, 2009. doi: https://doi.org/10.1090/S0273-0979-09-01249-X.

Eduardo Castro, Manel Martínez-Ramón, Godfrey Pearlson, Jing Sui, and Vince D. Calhoun. Characterization of groups using composite kernels and multi-source fmri analysis data: Application to schizophrenia. NeuroImage, 58(2):526 – 536, 2011.

F. Chazal and B. Michel. An introduction to topological data analysis: fundamental and practical aspects for data scientists, 2017.

Hongyoon Choi, Yu Kim, Hyejin Kang, Lee Hyekyoung, Hyung-Jun Im, Do Hwang, E Kim, June-Key Chung, and Dong Lee. Abnormal metabolic connectivity in the pilocarpine-induced epilepsy rat model: A multiscale network analysis based on persistent homology. NeuroImage, 99, 05 2014. doi: 10.1016/j.neuroimage.2014.05.039.

S. Chowdhury and F. Mémoli. Persistent Path Homology of Directed Networks, pages 1152–1169. 01 2018. ISBN 978-1-61197-503-1. doi: 10.1137/1.9781611975031.75.

M.K. Chung, J.L. Hanson, J. Ye, R.J. Davidson, and S.D. Pollak. Persistent homology in sparse regression and its application to brain morphometry. IEEE Transactions on Medical Imaging, 34, 08 2014. doi: 10.1109/TMI.2015.2416271.

D. Cohen-Steiner, H. Edelsbrunner, and J. Harer. Stability of persistence diagrams. Discrete Comput. Geom., 37:130–120, 2007.

Tamal Krishna Dey, Sayan Mandal, and William Varcho. Improved Image Classification using Topological Persistence. In Matthias Hullin, Reinhard Klein, Thomas Schultz, and Angela Yao, editors, Vision, Modeling and Visualization. The Eurographics Association, 2017. ISBN 978-3-03868-049-9. doi: 10.2312/vmv.20171272.

Olga Dunaeva, Herbert Edelsbrunner, Anton Lukyanov, Michael Machin, Daria Malkova, Roman Kuvaev, and Sergey Kashin. The classification of endoscopy images with persistent homology. Pattern Recognition Letters, 83:13–22, 2016. ISSN 0167-8655. doi: https://doi.org/10.1016/j.patrec.2015.12.012. Geometric, topological and harmonic trends to image processing.

H. Edelsbrunner, D. Letscher, and A. Zomorodian. Topological persistence and simplification. Discrete Comput. Geom., 28:511–533, 2002.

Cameron T. Ellis, Michael Lesnick, Gregory HenselmanPetrusek, Bryn Keller, and Jonathan D. Cohen. Feasibility of topological data analysis for event-related fmri. Network Neuroscience, 3(3):695–706, 2019. doi: 10.1162/netn\_a\_00095.

Karl J. Friston. Functional and effective connectivity in neuroimaging: A synthesis. Human Brain Mapping, 2(1-2): 56–78, 1994. doi: 10.1002/hbm.460020107.

Karl J. Friston. Functional and effective connectivity: A review. Brain Connectivity, 1(1):13–36, 2011. doi: 10.1089/brain.2011.0008. URL https://doi.org/10.1089/brain.2011.0008. PMID: 22432952.

Patrizio Frosini. A distance for similarity classes of submanifolds of a euclidean space. Bulletin of the Australian Mathematical Society, 42(3):407–415, 1990. doi: 10.1017/S0004972700028574.

Lazaros K. Gallos Hernán, A. Makse, and Mariano Sigman. A small world of weak ties provides optimal global integration of self-similar modules in functional brain networks. Proceedings of the National Academy of Sciences, 109(8):2825–2830, 2012. ISSN 0027-8424. doi: 10.1073/pnas.1106612109. URL https://www.pnas.org/content/109/8/2825.

Amanmeet Garg, Donghuan Lu, Karteek Popuri, and Mirza Faisal Beg. Cortical geometry network and topology markers for parkinson’s disease. ArXiv, 2016.

Noah Giansiracusa, Robert Giansiracusa, and Chul Moon. Persistent homology machine learning for fingerprint classification. pages 1219–1226, 12 2019. doi: 10.1109/ICMLA.2019.00201.

M. Gidea. Topology data analysis of critical transitions in financial networks. SSRN Electronic Journal, 01 2017. doi: 10.2139/ssrn.2903278.

Zeus Gracia-Tabuenca, Juan Carlos Díaz-Patiño, Isaac Arelio, and Sarael Alcauter. Topological data analysis reveals robust alterations in the whole-brain and frontal lobe functional connectomes in attention-deficit/hyperactivity disorder. bioRxiv, 2019. doi: 10.1101/751008.

D. Hartman, J. Hlinka, M. Paluš, D. Mantini, and M. Corbetta. The role of nonlinearity in computing graphtheoretical properties of resting-state functional magnetic resonance imaging brain networks. Chaos: An Interdisciplinary Journal of Nonlinear Science, 21(1):013119, 2011. doi: 10.1063/1.3553181. URL https://doi.org/10.1063/1.3553181.

A. Hatcher. Algebraic topology. Cambridge Univ. Press, Cambridge, 2000.

Jaroslav Hlinka, Milan Palus, Martin Vejmelka, Dante Mantini, and Maurizio Corbetta. Functional connectivity in resting-state fmri: Is linear correlation sufficient? NeuroImage, 54:2218–25, 02 2011. doi: 10.1016/j.neuroimage.2010.08.042.

Jaroslav Hlinka, David Hartman, Martin Vejmelka, Jakob Runge, Norbert Marwan, Juergen Kurths, and Milan Paluš. Reliability of inference of directed climate networks using conditional mutual information. Entropy, 15 (6):2023–2045, 2013. doi: 10.3390/e15062023.

D. Horak, S. Maletić, and M. Rajković. Persistent homology of complex networks. Journal of Statistical Mechanics: Theory and Experiment, (03), 2009.

A. Khalid, B.S. Kim, M.K. Chung, J.C. Ye, and D. Jeon. Tracing the evolution of multi-scale functional networks in a mouse model of depression using persistent brain network homology. NeuroImage, 101, 07 2014. doi: 10.1016/j.neuroimage.2014.07.040.

Jakub Kopal, Anna Pidnebesna, David Tomeček, Jaroslav Tintěra, and Jaroslav Hlinka. Typicality of Functional Connectivity robustly captures motion artifacts in rsfMRI across datasets, atlases and preprocessing pipelines. Human Brain Mapping, Submitted, 2020.

Jakub Kořenek and Jaroslav Hlinka. Causal network discovery by iterative conditioning: Comparison of algorithms. Chaos: An Interdisciplinary Journal of Nonlinear Science, 30(1):013117, 2020. doi: 10.1063/1.5115267.

Liqun Kuang, Xie Han, Caselli Chen, Kewei Reiman Richard J., and Yalin Eric M., Wang. A concise and persistent feature to study brain resting-state network dynamics: Findings from the alzheimer’s disease neuroimaging initiative. Human brain mapping, 40(4):1062–1081, 03 2019a. doi: 10.1002/hbm.24383.

Liqun Kuang, Deyu Zhao, Jiacheng Xing, Zhongyu Chen, Fengguang Xiong, and Xie Han. Metabolic brain network analysis of fdg-pet in alzheimer’s disease using kernel-based persistent features. Molecules, 24:2301, 06 2019b. doi: 10.3390/molecules24122301.

Dong Lee. Clinical personal connectomics using hybrid pet/mri. Nuclear Medicine and Molecular Imaging, 53:1–11, 01 2019. doi: 10.1007/s13139-019-00572-3.

H. Lee, M. K. Chung, H. Kang, B.-N. Kim, and D. S. Lee. Computing the shape of brain networks using graph filtration and gromov-hausdorff metric. In Gabor Fichtinger, Anne Martel, and Terry Peters, editors, Medical Image Computing and Computer-Assisted Intervention – MIC-CAI 2011, pages 302–309, Berlin, Heidelberg, 2011a. Springer Berlin Heidelberg. ISBN 978-3-642-23629-7.

H. Lee, M.K. Chung, H. Kang, B.-N. Kim, and D.S. Lee. Discriminative persistent homology of brain networks. Proceedings - International Symposium on Biomedical Imaging, pages 841–844, 03 2011b. doi: 10.1109/ISBI.2011.5872535.

H. Lee, H. Kang, M.K. Chung, B. Kim, and D.S. Lee. Persistent brain network homology from the perspective of dendrogram. IEEE Transactions on Medical Imaging, 31 (12):2267–2277, 12 2012.

Wei Liao, Zhiqiang Zhang, Zhengyong Pan, Dante Mantini, Jurong Ding, Xujun Duan, Cheng Luo, Guangming Lu, and Huafu Chen. Altered functional connectivity and small-world in mesial temporal lobe epilepsy. PloS one, 5 (1), 2010.

H. Lutkepohl. New Introduction to Multiple Time Series Analysis. Springer Publishing Company, Incorporated, 2007. ISBN 3540262393.

Mary-Ellen Lynall, Danielle S. Bassett, Robert Kerwin, Peter J. McKenna, Manfred Kitzbichler, Ulrich Muller, and Ed Bullmore. Functional connectivity and brain networks in schizophrenia. Journal of Neuroscience, 30(28):9477–9487, 2010. ISSN 0270-6474. doi: 10.1523/JNEUROSCI.0333-10.2010.

Daniel Lütgehetmann, Dejan Govc, Jason P. Smith, and Ran Levi. Computing persistent homology of directed flag complexes. Algorithms, 13(1):19, Jan 2020. ISSN 1999-4893. doi: 10.3390/a13010019. URL http://dx.doi.org/10.3390/a13010019.

P. Masulli and A. Villa. The topology of the directed clique complex as a network invariant. SpringerPlus, 5, 2016.

Emanuela Merelli, Marco Piangerelli, Matteo Rucco, and Daniele Toller. A topological approach for multivariate time series characterization: the epileptic brain. EAI Endorsed Transactions on Self-Adaptive Systems, 2(7), 5 2016. doi: 10.4108/eai.3-12-2015.2262525.

Pavol Mikolas, Jaroslav Hlinka, Antonin Skoch, Zbynek Pitra, Thomas Frodl, Filip Spaniel, and Tomas Hajek. Machine learning classification of first-episode schizophrenia spectrum disorders and controls using whole brain white matter fractional anisotropy. BMC Psychiatry, 18 (1), 2018.

J. R. Munkres. Elements of Algebraic Topology. Addison Wesley Publishing Company, Inc., 2725 Sand Hill Road Menlo Park, California 94025, 1984.

Isaura Oliver, Jaroslav Hlinka, Jakub Kopal, Jakob Runge, and Jörn Davidsen. Quantifying the variability in restingstate networks. Entropy, 21(9):882, 2019. doi: 10.3390/e21090882.

N. Otter, M. A. Porter, U. Tillmann, P. Grindrod, and H. A. Harrington. A roadmap for the computation of persistent homology. EPJ Data Science, 6(17), 2017.

D. Pachauri, C. Hinrichs, M.K. Chung, S.C. Johnson, and V. Singh. Topology-based kernels with application to inference problems in alzheimer’s disease. IEEE Transactions on Medical Imaging, 30:1760–70, 04 2011. doi: 10.1109/TMI.2011.2147327.

H.-J. Park and Karl Friston. Structural and functional brain networks: From connections to cognition. Science, 342 (6158):1238411–1238411, 2013. ISSN 0036-8075. doi: 10.1126/science.1238411.

G. Petri, M. Scolamiero, I. Donato, and F. Vaccarino. Topological strata of weighted complex networks. PLOS ONE, 8(6):1–8, 2013. doi: 10.1371/journal.pone.0066506. URL https://doi.org/10.1371/journal.pone.0066506.

A. Phinyomark, E. Ibáñez-Marcelo, and G. Petri. Restingstate fmri functional connectivity: Big data preprocessing pipelines and topological data analysis. IEEE Transactions on Big Data, 3(4):415–428, 2017. ISSN 2372-2096. doi: 10.1109/TBDATA.2017.2734883.

Marco Piangerelli, Matteo Rucco, Luca Tesei, and Emanuela Merelli. Topological classifier for detecting the emergence of epileptic seizures. BMC Research Notes, 11, 06 2018. doi: 10.1186/s13104-018-3482-7.

Archit Rathore, Sourabh Palande, Jeffrey Anderson, Brandon Zielinski, P. Fletcher, and Bei Wang. Autism Classification Using Topological Features and Deep Learning: A Cautionary Tale, pages 736–744. 10 2019. ISBN 978-3-030-32247-2. doi: 10.1007/978-3-030-32248-982.

M. W. Reimann, M. Nolte, M. Scolamiero, K. Turner, R. Perin, G. Chindemi, P. D-lotko, R. Levi, K. Hess, and H. Markram. Cliques of neurons bound into cavities provide a missing link between structure and function. Frontiers in Computational Neuroscience, 11:48, 2017. ISSN 1662-5188. doi: 10.3389/fncom.2017.00048.

Jan Reininghaus, Stefan Huber, Ulrich Bauer, and Roland Kwitt. A stable multi-scale kernel for topological machine learning. pages 4741–4748. Boston, MA, 06 2015. doi: 10.1109/CVPR.2015.7299106.

Harry Rubin-Falcone, Francesca Zanderigo, Binod Thapa-Chhetry, Martin Lan, Jeffrey M. Miller, M. Elizabeth Sublette, Maria A. Oquendo, David J. Hellerstein, Patrick J. McGrath, Johnathan W. Stewart, and J. John Mann. Pattern recognition of magnetic resonance imaging-based gray matter volume measurements classifies bipolar disorder and major depressive disorder. Journal of Affective Disorders, 227:498–505, 2018.

Matteo Rucco, Filippo Castiglione, Emanuela Merelli, and Marco Pettini. Characterisation of the idiotypic immune network through persistent entropy, pages 117–128. Proceedings of ECCS 2014. Lucca, Italy, September 2014. doi: 10.1007/978-3-319-29228-1\_11.

Mohammad Khubeb Siddiqui, Ruben Morales-Menendez, Xiaodi Huang, and Nasir Hussain. A review of epileptic seizure detection using machine learning classifiers. 7 (1):5, 2020. doi: 10.1186/s40708-020-00105-1.

Stephen M. Smith. The future of fmri connectivity. NeuroImage, 62(2):1257–1266, 2012. ISSN 1053-8119. doi: https://doi.org/10.1016/j.neuroimage.2012.01.022. URL http://www.sciencedirect.com/science/article/pii/S1053811912000390.20YEARSOFfMRI.

V. Solo, J. Poline, M. A. Lindquist, S. L. Simpson, F. D. Bowman, M. K. Chung, and B. Cassidy. Connectivity in fmri: Blind spots and breakthroughs. IEEE Transactions on Medical Imaging, 37(7):1537–1550, 2018.

Bernadette J. Stolz, Tegan Emerson, Satu Nahkuri, Mason A. Porter, and Heather A. Harrington. Topological Data Analysis of Task-Based fMRI Data from Experiments on Schizophrenia. arXiv e-prints, art. 1809.08504, 2018.

Guillaume Tauzin, Umberto Lupo, Lewis Tunstall, Julian Burella Pérez, Matteo Caorsi, Anibal MedinaMardones, Alberto Dassatti, and Kathryn Hess. giottotda: A topological data analysis toolkit for machine learning and data exploration, 2020.

Qawi Telesford, Sean Simpson, Jonathan Burdette, Satoru Hayasaka, and Paul Laurienti. The brain as a complex system: Using network science as a tool for understanding the brain. Brain connectivity, 1:295–308, 10 2011. doi: 10.1089/brain.2011.0055.

Christopher Tralie, Nathaniel Saul, and Rann Bar-On. Ripser.py: A lean persistent homology library for python. The Journal of Open Source Software, 3(29):925, Sep 2018. doi: 10.21105/joss.00925. URL https://doi.org/10.21105/joss.00925.

K. Turner. Rips filtrations for quasimetric spaces and asymmetric functions with stability results. Algebr. Geom. Topol., 19(3):1135–1170, 2019. doi: 10.2140/agt.2019.19.1135.

Pedro A. Valdes-Sosa, Alard Roebroeck, Jean Daunizeau, and Karl Friston. Effective connectivity: Influence, causality and biophysical modeling. NeuroImage, 58(2):339–361, 2011. ISSN 1053-8119. doi: https://doi.org/10.1016/j.neuroimage.2011.03.058. URL http://www.sciencedirect.com/science/article/pii/S1053811911003375.

Yuan Wang, Hernando Ombao, and Moo Chung. Topological seizure origin detection in electroencephalographic signals. Proceedings. IEEE International Symposium on Biomedical Imaging, 2015:351–354, 04 2015. doi: 10.1109/ISBI.2015.7163885.

Larry Wasserman. Topological data analysis. Annual Review of Statistics and Its Application, 5(1):501–532, 2018. doi: 10.1146/annurev-statistics-031017-100045.

Eleanor Wong, Sourabh Palande, Bei Wang, Brandon Zielinski, Jeffrey Anderson, and P. Fletcher. Kernel partial least squares regression for relating functional brain network topology to clinical measures of behavior. IEEE 13th International Symposium on Biomedical Imaging (ISBI), pages 1303–1306, 04 2016. doi: 10.1109/ISBI.2016.7493506.

Melih C. Yesilli, Firas A. Khasawneh, and Andreas Otto. Topological Feature Vectors for Chatter Detection in Turning Processes. arXiv e-prints, art. 1905.08671, 2019.

A. Zomorodian and G. Carlsson. Computing persistent homology. Discrete Comput. Geom., 33:249–274, 2005.

Mariana Zurita, Cristian Montalba, Tomás Labbé, Juan Pablo Cruz Josué, Dalboni da Rocha, Cristián Tejos, Ethel Ciampi, Claudia Cárcamo, Ranganatha Sitaram, and Sergio Uribe. Characterization of relapsingremitting multiple sclerosis patients using support vector machine classifications of functional and diffusion MRI data. 20:724–730. ISSN 2213-1582. doi: 10.1016/j.nicl.2018.09.002.

